# Quantifying quantify 64 drugs, illicit substances, and D- and L- isomers in human oral fluid with liquid-liquid extraction

**DOI:** 10.1101/2022.10.29.514362

**Authors:** Brian Robbins, Rob E. Carpenter, Mary Long, Jacob Perry

**Author notes:** Corresponding Author Rob E. Carpenter, PhD 10935 CR 159, Tyler, Texas 75703.

## Abstract

Although human oral fluid has become more routine for quantitative drug detection in pain management, detecting a large scope of medications and substances is costly and technically challenging for laboratories. This paper presents a quantitative assay for 64 pain medications, illicit substances, and drug metabolites in human oral fluid. The novelty of this assay is that it was developed on an older model AB SCIEX 4000 instrument and renders obscure the need for more technical and expensive laboratory equipment. This method includes addition of internal standard and a 2-step liquid-liquid extraction and dry-down step to concentrate and clean the samples. The samples were suspended in 50% MeOH in water and separation and detection was accomplished using triple quadrupole mass spectrometry (LC-MS/MS). Separation was achieved using reverse-phase liquid chromatography with detection by LC-MS/MS. A second injection was done in negative mode to determine THC-COOH concentration as an indicator of THC. An aliquot of the (already) extracted samples was analyzed for D- and L- isomers of amphetamine and methamphetamine using a chiral column. The standard curve spanned from 5 to 2000 ng/mL for most of the analytes (1 to 2000 ng/mL for fentanyl and THC-COOH) and up to 1000 ng/mL for 13 analytes. Pregabalin and gabapentin ranged from 25 to 2000 ng/mL. The result is a low-cost method for the sensitive detection of a wide-ranging oral fluid menu for pain management. This assay has a high sensitivity, and good precision and accuracy for all analytes with an older model mass spectrometer.

## 1 INTRODUCTION

The United States Controlled Substances Act (CSA) places drugs and certain chemicals used to make drugs into five distinct categories or schedules depending upon the drug’s acceptable medical use and its potential for abuse or dependency.^1^ The abuse rate is a determinate factor in the scheduling of the drug; for example, Schedule I drugs are considered the most dangerous class of drugs with a high potential for abuse and potentially severe psychological and/or physical dependence. As the drug schedule changes (Schedule II, Schedule III, etc.), so does the abuse potential—Schedule V drugs represents the least potential for abuse.

Scheduled drugs are some of the most powerful tools used in the treatment of chronic pain, anxiety, depression, and attention deficit disorders. For example, opiates are commonly prescribed for the management of acute or chronic pain despite research that long term biological efficacy is questionable.^2^ To assist in the management of chronic pain, clinicians have opted for testing patients for compliance to their drug regimen. Routine assessment for non-compliance or non-medical use is frequently accomplished through urine drug testing (UDT) based on risk of drug misuse, abuse, and diversion. Although UDT is considered the common practice for detecting scheduled drug compliance, often patients are unable to provide a urine sample for various reasons. In this case, oral fluid drug testing (ODT) can serve as an effective alternative to UDT for medication monitoring.^3^

Oral fluid drug testing is increasingly emerging as an alternative biological matrix for detecting drugs and monitoring patient medication compliance.^4–6^ Moreover, in certain clinical situations clinicians may find oral fluid more beneficial for detection of specific drugs over UDT.^7^ The matrix allows for easy collection, but attention to recovery, stability, and dilutions issues of some collection devices should be given consideration for pharmacokinetic studies.^7, 8^ Although ODT opioid assays that use dilute-and-shoot methods with little sample manipulation have been developed and validated on AB SCIEX 4500 instruments with excellent calibration ranges (2.5-1,000 ng/mL)^9^ robust ODT assays that also quantify the dextro (D-) and levo (L-) isomers of amphetamine and methamphetamine are less common. Furthermore, large ODT assays (50 drugs) have been successfully developed and validated on ultra-high performance triple quadrapole mass spectrometer (LC-MS/MS),^10^ yet large ODT assay development on older LC-MS/MS technology and separation specificity are less common. This work highlights the development and validation of a fast, accurate, and inexpensive method to quantify 64 medications, illicit substances, and the D- and L- isomers of amphetamine and methamphetamine in human oral fluid specimens using liquid-liquid extraction and LC-MS/MS with an older model AB SCIEX 4000 instrument. Each sample was initially analyzed for 63 targeted analytes using LC-MS/MS and the same extracts injected a second time using a delta-9 tetrahydrocannabinol (THC) specific electrospray (ES) negative assay to detect the THC metabolite 11-Nor-9-carboxy-Δ9-tetrahydrocannabinol (THC-COOH). Then, for samples that initially showed positive for amphetamine or methamphetamine an additional sample was taken from the already extracted specimen and analyzed with a newly developed assay designed specifically to determine the D- and L- isomer status to define non-illicit versus illicit etiology. Accordingly, this work presents the development and validation of a robust human oral fluid drug assay—referred to in this paper as P63 assay. The assay development and validation are offered here for the benefit of high-throughput laboratories that seek novel solutions for a scoping ODT menu with fast and accurate chemical analysis using less expensive older model AB SCIEX instruments.

## 2 METHODS AND MATERIALS

### 2.1 Reagents and standards

All analyte stock solutions at 1 mg/mL concentrations and deuterated internal standards at 100 µg/mL were purchased from Cerilliant Corporation (Round Rock, TX, USA). All organic solvents including methanol, acetonitrile, formic acid (88%), dichloromethane, 2 propanol and ethyl acetate were obtained from Fisher Scientific (Pittsburgh, PA, USA). Oral fluid Quantisal® extraction buffer and collection devices were obtained from Immunalysis Corporation (Pomona, CA, USA).

### 2.2 Mobile phase and extraction solutions

Mobile phase A (MPA) solution was created with water, methanol, and formic acid (97.4:2.5:0.1) by using a 1L bottle to combine 974 mL of LC/MS grade water and 25 mL of LC/MS grade methanol; then 1 mL of formic acid (88%) was added and mixed thoroughly. The solution was stored at room temperature for up to two weeks. Mobile phase B (MPB) solution was created with acetonitrile and methanol (1:1) by using a 1L bottle to combine 500 mL of LC/MS-grade methanol and 500 mL of LC/MS-grade acetonitrile. This solution can be kept at room temperature for up to 1 year. A D- and L- mobile phase (MPDL) solution was created by adding ∼993.2 mL of methanol to a 1L bottle Then, using a mechanical pipette, 5 mL of type I water, 1.5 mL of acetic acid, and 0.3 mL of ammonium hydroxide were added and mixed thoroughly. This solution can be kept at room temperature for up to 1 year. A needle wash solution was created with methanol, acetonitrile, and clinical grade water (1:1:1) using a 1L bottle by adding equal volumes of methanol, acetonitrile, and water. Extraction solution 1 (ES1) was created with 50% dichloromethane and 50% 2-propanol by using a graduated cylinder under a fume hood. Equal volumes of dichloromethane and 2-propanol were added to a clean reagent bottle which was capped and mixed well. Extraction solution 2 (ES2) was created with 50% dichloromethane and 50% ethyl acetate by using a graduated cylinder under a fume hood. Equal volumes of dichloromethane and ethyl acetate were added to a clean reagent bottle which was capped and mixed well.

### 2.3 Standard preparation

An 8000 ng/mL stock solution was made by combining analyte stock controls and diluting it with MPA. In contrast, D- and L- amphetamine and methamphetamine were added in an amount to make a 4000 ng/mL stock of each isomer so that combined they would produce an 8000 ng/mL solution of total amphetamine and methamphetamine. In contrast, this means that the range of the d and l SC is from 2.5 to 1000 ng/mL (half the concentration). The resulting stock standard was diluted with MPA to produce the standard curve (SC). Concentrations were 8000 (undiluted), 4000, 2000, 1000 400, 200, 100, 40, 20, 10, 4 and 2 ng/mL. These solutions were stored at the concentrations above. They underwent a dilution during the assay (1 part standard to 3 parts mobile phase and THC standard) to achieve the concentration desired in sample analysis with oral fluid (saliva). The standards and quality control (QC) were diluted (0.5 mL) with 1.5 mL of extraction buffer. This approximates the condition seen with saliva after collection with the Quantisal® oral fluid sample collection device. The final concentration in the 0.5 mL sample SC included the following points: 2000, 1000, 500, 250, 100, 50, 25, 10, 5, 2.5, 1 and 0.5 ng/mL.

The assay QCs were made similarly; first making a 7200 ng/mL spiking solution in MPA then diluting to 3200, 2400, 300, 60, 12, and 2 ng/mL. The D- and L- amphetamine and methamphetamine QCs were made at half concentrations. Final concentrations of each QC were 1800, 800, 600, 75, 15, 3 and 0.5 ng/mL, after the 1:4 dilution with MPA and THC QC same as the SC points noted above.

The internal standard working solution (ISWS) for the P63 assay was made by filling a 100 mL graduated cylinder to the 50 mL mark with 10% methanol in water and adding 250 µL of each of the internal standards listed above. The volume was brought to 100 mL with additional 10% methanol producing a concentration of 250 ng/mL.

The 20000 ng/mL THC-COOH analyte stock solution was made by adding 200 µL of 1.0 mg/mL THC-COOH stock to a 15 mL polypropylene tube and bringing the volume to 10.0 mL using 50 % methanol in water. This was diluted further with 50% methanol in water to produce a SC with concentrations of 20000, 10000, 5000, 2500, 1000, 500, 250, 100, 50, 25, 10 and 5 ng/mL. Each solutions underwent a dilution during the assay that resulted in final concentrations of 2000, 1000, 500, 250, 100, 50, 25, 10, 5, 2.5, 1 and 0.5 ng/mL.

The THC-COOH 18000 ng/mL quality control stock solution was made by adding 180 µL of 1.0 mg/mL THC-COOH stock to a 15 mL polypropylene tube and bringing the volume to 10.0 mL using 50 % methanol in water. This was diluted further with 50% methanol in water to produce a SC with concentrations of 18000, 8000, 6000, 750, 150, 30 and 5 ng/mL. These solutions underwent a dilution during the assay that results in final concentrations of 1800, 800, 600, 75, 15, 3 and 0.5 ng/mL.

The THC-COOH ISWS was made by diluting 300 µL of 100 µg/mL THC-COOH-D9 (Cerilliant) internal standard with 40 mL of 0.1 M ammonium acetate buffer pH 5 made by weighing out 7.7g of LC-MS/MS grade ammonium acetate and transferring to a 1-liter bottle, add ∼700 mL of medical grade water that was capped and mixed until dissolved. The pH was adjusted to 5.0 with glacial acetic acid (3 shots of 733 µl) QS to 1L with medical grade water then capped and mixed well and stored at 2-8 C for up to 2 months.

### 2.4 Instrumentation

The liquid chromatography components of the LC-MS/MS system consisted of a model CBM- 20A controller, 2 model Prominence LC-20AD pumps, a model DGU-20A5 degasser and a model SIL-20AC autosampler all obtained from (Shimadzu, Columbia MD, USA, based in Kyoto, Japan). The mass spectrometer used was a SCIEX API 4000 and the acquisition software was Analyst, v 1.5.2, build 5704 (Framingham, MA, USA). Nitrogen was obtained using a Peak ABN2ZA gas generator (Peak Scientific, Billerica, MA, USA). Reagents were weighed on a Mettler Toledo MX5 analytical micro balance (Fisher Scientific, Pittsburgh, PA, USA). Samples were dried on a TurboVap*®* LV (Uppsala, Sweden). Samples were vortexed on a Fisherbrand 120 multitube vortex. The analytical column was a Restek Ultra Biphenyl 5.0 µm (2.1 x 50 mm column), Catalog # 9109552 and the guard column was a Restek Ultra Biphenyl 5.0 µm (10 x 4 mm column) Catalog # 910950210 (Restek, Bellefonte, PA, USA) and Astec CHIROBIOTIC® V2 5.0 µm (2.1mm x 25 cm column) Catalog # 15020AST SUPLECO®, (Bellefonte, PA, USA).

### 2.5 Analyte optimization

Individual analytes and internal standards were optimized by using T-infusion with 50% MPB mobile phase and tuning for declustering potential (DP), entrance potential (EP), collision energy (CE) and exit potential (CXP) at a flow rate of 0.7 mL/min. The two most abundant fragments were selected for monitoring using MRM. This resulted in the settings presented in Appendix A.

### 2.6 Sample preparation and procedures

Standards and QC were prepared in 13 x 100 mm tubes by adding 125 µl of P63 standard or QC, 50 µl of THC-COOH standard or QC, and 325 µl of MPA. A volume of 1.5 mL of Quantisal® buffer was added to all SC and QC preparations to match the volume of sample that is routinely extracted. A 2 mL aliquot of patient sample collected with the Quantisal® device was extracted along with the standards and QC in a vented biological safety cabinet and transferred to a properly labeled 13×100 mm glass tube. The ISWS for the P63 and THC-COOH assays were added to all tubes except for double blank and wash tubes. The samples were extracted with 2 mL of 50% dichloromethane:50% 2 propanol. The samples were mixed with a mass vortex for 5 min. The tubes were transferred to centrifuge buckets, covered, and centrifuged 10 min at 3000 rpm (1690 x g) to facilitate separation. The clear bottom layer was transferred to a fresh 13 x 100 mm borosilicate glass tube. The remaining blue aqueous phase was extracted further with 2 mL of 50% dichloromethane and 50% ethyl acetate and mixed with a mass vortex for 5 min. The samples were centrifuged an additional 10 min at 3000 rpm (1690 x g) to fully separate the resultant layers. The bottom clear layer was combined with the corresponding clear aliquot from the first extraction step. The clear organic samples were dried under nitrogen in a TurboVap*®* LV.

Sample preparation for the LC-MS/MS was done by adding 125 µL of 50% MeOH and 50% water to each sample, SC and QC tube, wait for 10-15 min to dissolve, then vortex. Then 100 µL of the re-suspended sample was transferred to a corresponding well on the preparation plate. Next 400 µL of MPA was added to each well, the plate was covered with a plate mat and mixed briefly. The plate was centrifuged for 15 min at 4000 rpm (2272 x g) and samples were transferred (∼300 µL) to a fresh, labeled plate. It was covered with a plate mat and analyzed for P63 and then THC-COOH.

Sample preparation for D- and L- analysis by LC-MS/MS involved transferring 50 µL of the already extracted standards, QC, and any samples of interest to a new plate. Then 450 µL of MPDL was added to each well and mixed with a multichannel pipette, the plate was covered with a plate mat and analyzed for the D- and L- isomers of amphetamine and methamphetamine.

The LC-MS/MS conditions and separation parameters for all three methods/assays used in this study are expanded in Tables 1-3. In summary samples were injected with the P63 parameters followed by a second injection with the THC-COOH parameters. A 50 µL aliquot of these prepared samples, standards and quality controls were transferred to a fresh plate and diluted with 450 µL of D- and L- assay mobile phase for LC-MS/MS injection to determine D- and L- isomer concentrations.

**TABLE 1.** Gradient for separation of 63 ES positive analytes consisting of a series of linear step gradients over 8 min.

| Time<br>(minutes) | Flow<br>(mL/min) | % MPA | % MPB | Curve |
| --- | --- | --- | --- | --- |
| 0 | 0.7 | 93 | 7 | Linear |
| 0.1 | 0.7 | 93 | 7 | Linear |
| 3 | 0.7 | 60 | 40 | Linear |
| 4.5 | 0.7 | 10 | 90 | Linear |
| 6 | 0.7 | 10 | 90 | Linear |
| 6.1 | 0.7 | 93 | 7 | Linear |
| 7.6 | 0.7 | 93 | 7 | Linear |
| 8.02 | Stop |  |  |  |

*Note:* electrospray (ES); mobile phase A (MPA); mobile phase B (MPB)

**TABLE 2.**
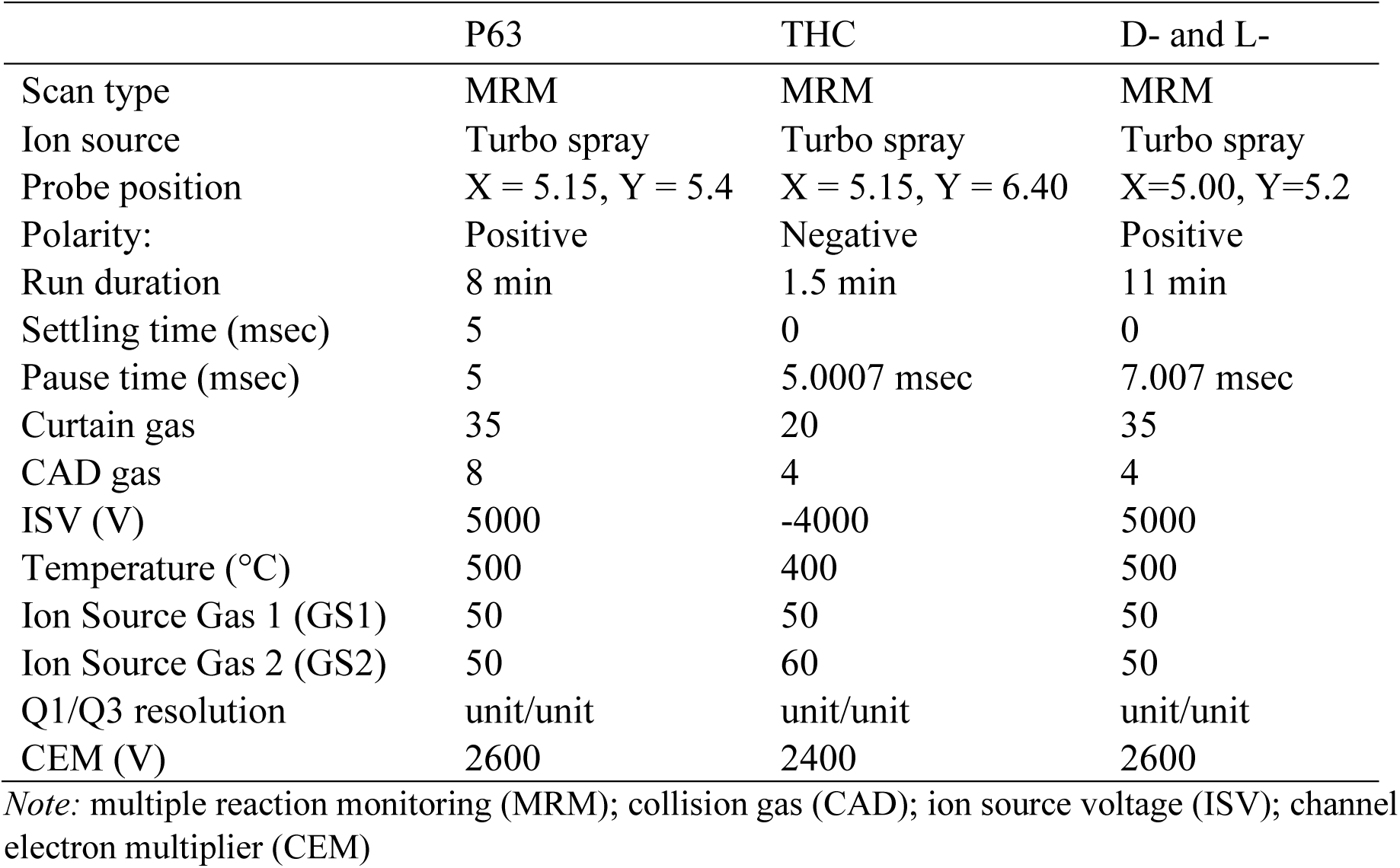
LC-MS/MS conditions for the three assays on a single oral fluid sample.

|  | P63 | THC | D- and L- |
| --- | --- | --- | --- |
| Scan type | MRM | MRM | MRM |
| Ion source | Turbo spray | Turbo spray | Turbo spray |
| Probe position | X = 5.15, Y = 5.4 | X = 5.15, Y = 6.40 | X=5.00, Y=5.2 |
| Polarity: | Positive | Negative | Positive |
| Run duration | 8 min | 1.5 min | 11 min |
| Settling time (msec) | 5 | 0 | 0 |
| Pause time (msec) | 5 | 5.0007 msec | 7.007 msec |
| Curtain gas | 35 | 20 | 35 |
| CAD gas | 8 | 4 | 4 |
| ISV (V) | 5000 | -4000 | 5000 |
| Temperature (°C) | 500 | 400 | 500 |
| Ion Source Gas 1 (GS1) | 50 | 50 | 50 |
| Ion Source Gas 2 (GS2) | 50 | 60 | 50 |
| Q1/Q3 resolution | unit/unit | unit/unit | unit/unit |
| CEM (V) | 2600 | 2400 | 2600 |

*Note:* multiple reaction monitoring (MRM); collision gas (CAD); ion source voltage (ISV); channel electron multiplier (CEM)

**TABLE 3.** Oral fluid inlet settings for P63, THC, and D- and L- panels

| Inlet Settings | P 63 and THC Panels | D and L Panel |
| --- | --- | --- |
| Analytical Column | Restek Ultra Biphenyl 50 x 2.1 mm, 5 μm | Supelco Astek Chirobiotic V 250 x 2.1 mm, 5 μm |
| Guard Cartridge | Restek Ultra Biphenyl 10 x 4.0 mm, 5 μm | None |
| Sample Temperature | 15 ± 5.0°C | 15 ± 5.0°C |
| Column Temperature | 30.0 ± 5.0°C | 30.0 ± 5.0°C |
| Mobile Phase A | Water:MeOH:Formic acid – 97.5:2.5:0.1 | Water:Acetic Acid:Ammonium Hydroxide: Methanol 5:1:0.3:993.5 |
| Mobile Phase B | MeOH:Acetonitrile – 1:1 | N/A |
| Needle Rinse | 1:1:1 | Water:Acetic Acid:Ammonium Hydroxide: Methanol 5:1:0.3:993.5 |
| Flow Rate | 0.7 mL/min | 0.3 mL/min |
| Injection Volume | 10 μL | 10 μL |
| Run Time | 8 min, 1.5 min | 11 min |

### 2.7 Method validation procedures

#### 2.7.1 Matrix lot-to-lot comparison

Individual lots of human matrix (saliva) differ according to a person’s overall health and hydration status.^11^ A single lot of oral fluid is not enough to demonstrate the ruggedness of the assay system when such variability in the matrix exists.^12^ Due to this, and in accordance with current CAP standards, a minimum of 10 lots of human matrix were collected from donors who verify that they are not taking the analytes that are being validated. These donor oral fluid samples were spiked at a low-level concentration with each analyte. These samples were prepared, extracted, and analyzed as described above. The responses were calculated and the analyte to internal standard (IS) ratio and %CV is shown in Table 10.

#### 2.7.2 Analytical measurement range

The analytical measurement range (AMR) of the assay refers to the concentration range that the assay is validated within and is determined by running a series of calibration curve standards covering a concentration range that encompass the concentration of analyte expected to find in patient samples.^13^ The limits of the AMR were bounded by the lower limit of quantitation (LLOQ) and the upper limit of quantitation (ULOQ). The dynamic range may be described by a linear or quadratic fit.^13, 14^ Calibration curves were created using a minimum of six non-zero calibration points. To be accepted as the AMR, all points describing the calibration curve must pass within ± 20% of the nominal concentration^13^. Furthermore, the correlation coefficient (R^2^) for the calibration curve must be ≥ 0.98, or R should be ≥0.99 to be acceptable.^15, 16^

#### 2.7.3 Sensitivity

The sensitivity of the assay system refers to the ability to reliably produce a signal throughout the entire calibration range, but specifically at the low-end of the calibration curve.^17^ In hyphenated mass spectrometry assays, a signal that produces a signal to noise ratio (S/N) of ≥10 is considered valid for the LLOQ of an assay system.^18^ Further, a S/N ratio of ≥5 is considered clear enough for the limit of detection. We test the sensitivity of the assay system by injecting 6 replicates of the LLOQ over 3 days and evaluating the resulting analytical determinations. Standard acceptance criteria of ±20% of nominal concentration apply.

#### 2.7.4 Intra-day precision and accuracy

Intra-day precision and accuracy were determined using six replicates of each of three quality control (QC) sample determinations and LLOQ from across at least three validation runs. Concentrations of the QC samples ranged across the curve, with the low QC set at approximately 3 times the LLOQ or less, the mid QC near the middle of the linear range and the high QC set at 80-90% of the ULOQ. Percent accuracy was determined for each individual measurement using the equation:

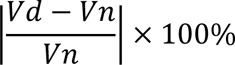

Where *Vd* is the concentration determined from the calibration curve and *Vn* is the nominal concentration for the QC standard. Precision was determined for each standard level by first determining the standard deviation of the six replicate standards and then applying the following equation:

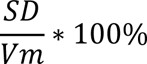

Where *SD* is the standard deviation of the six replicates and *Vm* is the mean value of the standard. To be accepted, the precision and accuracy for the replicate determinations must be ≤20% at each level.

#### 2.7.5 Inter-day precision and accuracy

Inter-day precision and accuracy were determined using all replicates of each of three quality control (QC low, QC mid, and QC high) and LLOQ sample determinations from the analytical runs performed on 3 separate days. Concentrations of the QC samples ranged across the curve, with the low QC set around 3 times the LLOQ, the mid QC near the middle of the linear range, and the high QC set at 80-90% of the ULOQ.

#### 2.7.6 Exogenous interfering substances

Drugs that are known or suspected of interfering with similar bioanalytical systems should be evaluated to ensure that they do not suppress ionization or cause false-positive results for a given analyte.^19, 20^ The following medications were evaluated: over-the-counter mix (consisting of acetaminophen, ibuprofen, pseudoephedrine, caffeine, and naproxen), salicylic acid, phenylephrine, phentermine, diphenhydramine, and dextromethorphan. A high concentration of the possible interfering drug (typically 2,000 ng/mL or greater) was spiked into a low QC sample (15 – 75 ng/mL low QC). Acceptance criteria for a substance to be deemed as non-interfering is that the quantitated value for the low QC should be within ± 20% of the nominal value.^21^ Furthermore, the spiked substance should not cause a false-positive or a false-negative result.

#### 2.7.7 Partial volumes and dilutions

A spiked solution was created at a concentration above the ULOQ in this case 4000 ng/mL. The sample was run at discrete dilutions 1:5, 1:10, 1:20, and 1:50. Concentration determinations for all dilutions should be within ± 20% of the nominal value following correction for the dilution factor.^21, 22^ More recent literature suggests that the signal to noise ratio of both the quantification trace and the qualifying ion trace be 3-10.^23, 24^ On occasion, an analyte will not have a qualifying ion that passes this criterion while still permitting the quantification trace to remain in a meaningful range. These instances should be documented in the laboratory SOP or validation report.

#### 2.7.8 Carryover

Carryover is the presence of an analyte in a blank injection following a positive injection, resulting in a false-positive sample.^25^ The injection needle should be washed in-between samples with a needle wash solution that is intended to remove contamination from the surface of the needle. The efficiency of this process is monitored during validation by assessing carryover in the following manner. Samples are injected in the following sequence: high QC, wash, high QC, wash, high QC, wash. Peak areas are integrated for both the analyte and internal standard. Peak area in the wash solutions should be 0.1 % or less of that found in the High QC standard. In addition, the mean of the peak area in the three wash solutions following the high QC replicates should be less than 20% of the LLOQ being used for the assay.^26^

## 3 RESULTS

### 3.1 Inter-day average back calculated calibration standards

Each validation run contained calibration standards with theoretical concentrations of 1, 2.5, 5, 10, 25, 50, 100, 500, 1000 and 2000 ng/mL of each of the analytes with an additional negative run at 0.5 ng/mL. These line up with the P63 standard concentrations for ease of addition. The calibration curves were determined by plotting the theoretical concentration versus the area ratio for each standard. A weighted (1/x; where x=concentration) quadratic regression line was fit to the data and used to determine the concentration of unknown samples. Table 4 shows the range of standard curves of the analytes and the correlation information. D- and L- curve concentrations were half the above concentration ranging from 0.25 (neg) to 1000 ng/mL. Mean R values were all at least 0.99 indicating good fit to the data.

**Table 4.** Statistical analysis for each analyte standard curve over three assays

| Drug / Metabolite | Curve Range (ng/mL) | Mean R | RSD | Mean Slope | SD Slope | N | Fit |
| --- | --- | --- | --- | --- | --- | --- | --- |
| 6-MAM | 5-2000 | 0.9988 | 0.0033 | 0.0340 | 0.0014 | 3.0000 | Quadratic |
| 7-Amino Clonazepam | 5-2000 | 0.9991 | 0.0006 | 0.0303 | 0.0129 | 3.0000 | Quadratic |
| Alprazolam | 5-2000 | 0.9996 | 0.0002 | 0.0762 | 0.0033 | 3.0000 | Quadratic |
| Amitriptyline | 5-1000 | 0.9989 | 0.0005 | 0.0106 | 0.0006 | 3.0000 | Quadratic |
| Amphetamine* | 5-1000 | 0.9990 | 0.0006 | 0.0594 | 0.0037 | 3.0000 | Quadratic |
| a-OH Alprazolam | 5-2000 | 0.9996 | 0.0003 | 0.0419 | 0.0028 | 3.0000 | Quadratic |
| Benzoylcegonine | 5-2000 | 0.9985 | 0.0012 | 0.0018 | 0.0001 | 3.0000 | Quadratic |
| Buprenorphine | 5-2000 | 0.9999 | 0.0001 | 0.0241 | 0.0007 | 3.0000 | Quadratic |
| Carisoprodol | 5-2000 | 0.9992 | 0.0006 | 0.0098 | 0.0010 | 3.0000 | Quadratic |
| Chlordiazepoxide* | 5-1000 | 0.9998 | 0.0000 | 0.0111 | 0.0003 | 3.0000 | Quadratic |
| Clonazepam | 5-2000 | 0.9991 | 0.0006 | 0.0161 | 0.0039 | 3.0000 | Quadratic |
| Codeine | 5-2000 | 0.9996 | 0.0005 | 0.0406 | 0.0064 | 3.0000 | Quadratic |
| Cotinine | 5-2000 | 0.9998 | 0.0001 | 0.0463 | 0.0001 | 3.0000 | Quadratic |
| Cyclobenzaprine | 5-2000 | 0.9999 | 0.0001 | 0.0054 | 0.0008 | 3.0000 | Quadratic |
| Desalkylflurazepam | 5-2000 | 0.9994 | 0.0001 | 0.0383 | 0.0010 | 3.0000 | Quadratic |
| Desipramine | 5-2000 | 0.9993 | 0.0006 | 0.0578 | 0.0047 | 3.0000 | Quadratic |
| Diazepam | 5-2000 | 0.9993 | 0.0005 | 0.1009 | 0.0117 | 3.0000 | Quadratic |
| Dihydrocodeine | 5-2000 | 0.9986 | 0.0008 | 0.0479 | 0.0028 | 3.0000 | Quadratic |
| Doxepin | 5-2000 | 0.9998 | 0.0001 | 0.0211 | 0.0006 | 3.0000 | Quadratic |
| EDDP | 5-2000 | 0.9997 | 0.0001 | 0.0106 | 0.0004 | 3.0000 | Quadratic |
| Fentanyl | 5-2000 | 0.9998 | 0.0001 | 0.0108 | 0.0006 | 3.0000 | Quadratic |
| Fluoxetine | 5-2000 | 0.9995 | 0.0004 | 0.0510 | 0.0028 | 3.0000 | Quadratic |
| Gabapentin | 25-2000 | 0.9961 | 0.0033 | 0.0609 | 0.0485 | 3.0000 | Quadratic |
| Hydrocodone | 5-2000 | 0.9992 | 0.0005 | 0.0123 | 0.0018 | 3.0000 | Quadratic |
| Hydromorphone | 5-2000 | 0.9983 | 0.0021 | 0.1583 | 0.0211 | 3.0000 | Quadratic |
| Imipramine | 5-2000 | 0.9999 | 0.0002 | 0.0067 | 0.0007 | 3.0000 | Quadratic |
| Ketamine* | 5-1000 | 0.9990 | 0.0009 | 0.0107 | 0.0015 | 3.0000 | Quadratic |
| Lorazepam | 5-2000 | 0.9995 | 0.0001 | 0.0091 | 0.0003 | 3.0000 | Quadratic |
| MDA* | 5-1000 | 0.9994 | 0.0004 | 0.0550 | 0.0005 | 3.0000 | Quadratic |
| MDMA | 5-2000 | 0.9998 | 0.0002 | 0.0050 | 0.0004 | 3.0000 | Quadratic |
| MDPV | 5-2000 | 0.9998 | 0.0001 | 0.0777 | 0.0019 | 3.0000 | Quadratic |
| Meperidine* | 5-1000 | 0.9995 | 0.0004 | 0.0315 | 0.0020 | 3.0000 | Quadratic |
| Meprobamate | 5-2000 | 0.9997 | 0.0003 | 0.0102 | 0.0002 | 3.0000 | Quadratic |
| Methadone | 5-2000 | 0.9986 | 0.0006 | 0.0080 | 0.0001 | 3.0000 | Quadratic |
| Methamphetamine | 5-2000 | 0.9999 | 0.0001 | 0.0104 | 0.0004 | 3.0000 | Quadratic |
| Methylphenidate* | 5-1000 | 0.9993 | 0.0002 | 0.0167 | 0.0007 | 3.0000 | Quadratic |
| Morphine | 5-2000 | 0.9988 | 0.0008 | 0.0538 | 0.0038 | 3.0000 | Quadratic |
| Naloxone | 5-2000 | 0.9994 | 0.0004 | 0.0548 | 0.0061 | 3.0000 | Quadratic |
| Naltrexone | 5-2000 | 0.9991 | 0.0013 | 0.1273 | 0.0067 | 3.0000 | Quadratic |
| Nicotine | 5-2000 | 0.9996 | 0.0002 | 0.0712 | 0.0033 | 3.0000 | Quadratic |
| Norbuprenorphine | 5-2000 | 0.9997 | 0.0002 | 0.0444 | 0.0026 | 3.0000 | Quadratic |
| Nordiazepam | 5-2000 | 0.9991 | 0.0008 | 0.0531 | 0.0061 | 3.0000 | Quadratic |
| Norfentanyl | 5-2000 | 0.9996 | 0.0002 | 0.0508 | 0.0057 | 3.0000 | Quadratic |
| Norhydrocodone | 5-2000 | 0.9993 | 0.0004 | 0.0858 | 0.0079 | 3.0000 | Quadratic |
| Norketamine | 5-2000 | 0.9998 | 0.0001 | 0.0037 | 0.0004 | 3.0000 | Quadratic |
| Noroxycodone | 5-2000 | 0.9990 | 0.0005 | 0.0032 | 0.0000 | 3.0000 | Quadratic |
| Noroxymorphone | 5-2000 | 0.9998 | 0.0001 | 0.0040 | 0.0002 | 3.0000 | Quadratic |
| Nortriptyline | 5-2000 | 0.9983 | 0.0007 | 0.0144 | 0.0006 | 3.0000 | Quadratic |
| O-desmethyl Tramadol | 5-2000 | 0.9996 | 0.0002 | 0.0226 | 0.0010 | 3.0000 | Quadratic |
| Oxazepam | 5-2000 | 0.9999 | 0.0002 | 0.0249 | 0.0075 | 3.0000 | Quadratic |
| Oxycodone | 5-2000 | 0.9992 | 0.0004 | 0.1193 | 0.0112 | 3.0000 | Quadratic |
| Oxymorphone | 5-2000 | 0.9989 | 0.0014 | 0.0092 | 0.0006 | 3.0000 | Quadratic |
| Paroxetine | 5-2000 | 0.9992 | 0.0003 | 0.0063 | 0.0001 | 3.0000 | Quadratic |
| Phencyclidine | 5-2000 | 0.9997 | 0.0001 | 0.0025 | 0.0002 | 3.0000 | Quadratic |
| Pregabalin | 25-2000 | 0.9982 | 0.0023 | 0.0784 | 0.0118 | 3.0000 | Quadratic |
| Ritalinic Acid* | 5-1000 | 0.9995 | 0.0005 | 0.0094 | 0.0000 | 3.0000 | Quadratic |
| Tapentadol | 5-2000 | 0.9994 | 0.0006 | 0.0085 | 0.0006 | 3.0000 | Quadratic |
| Temazepam | 5-2000 | 0.9998 | 0.0001 | 0.0189 | 0.0177 | 3.0000 | Quadratic |
| Tramadol* | 5-1000 | 0.9994 | 0.0004 | 0.0092 | 0.0010 | 3.0000 | Quadratic |
| Venlafaxine | 5-2000 | 0.9996 | 0.0004 | 0.0092 | 0.0010 | 3.0000 | Quadratic |
| Zaleplon | 5-2000 | 0.9996 | 0.0004 | 0.0460 | 0.0014 | 3.0000 | Quadratic |
| Zolpidem | 5-2000 | 0.9999 | 0.0001 | 0.0100 | 0.0004 | 3.0000 | Quadratic |
| Zopiclone | 5-2000 | 0.9991 | 0.0007 | 0.7457 | 1.1989 | 3.0000 | Quadratic |
| THC-COOH | 1-2000 | 0.9996 | 0.0004 | 0.0029 | 0.0015 | 3.0000 | Quadratic |
| D-Amphetamine | 2.5-1000 | 0.9994 | 0.0003 | 0.0104 | 0.0010 | 3.0000 | Quadratic |
| L-Amphetamine | 2.5-1000 | 0.9993 | 0.0006 | 0.0105 | 0.0010 | 3.0000 | Quadratic |
| D-Methamphetamine | 2.5-1000 | 0.9995 | 0.0008 | 0.0218 | 0.0021 | 3.0000 | Quadratic |
| L-Methamphetamine | 2.5-1000 | 0.9985 | 0.0021 | 0.0246 | 0.0031 | 3.0000 | Quadratic |

*Note:* \*Denotes ULOQ of 1000 ng/mL, Gabapentin and Pregabalin had a range from 25 to 2000 ng/mL.

### 3.2 Accuracy and precision, LLOQ

Six replicates of each validation level were run on at least 3 days. The theoretical concentrations were 1, 5 or 25 for LLOQ on P63, 3, 15 or 75 for QC low, 600 ng/mL for QC mid and 800 or 1800 for the QC high values. The THC-COOH assay had concentrations of 1 ng/mL for the LLOQ, 3 ng/mL for the QC low, 600 for the QC mid and 2000 for the QC high. The D- and L- assay had an LLOQ of 2.5 ng/mL with a QC low of 7.5 ng/mL, a QC mid of 300 ng/mL and a high QC of 1000 ng/mL. Tables 5 & 6 indicates inter-assay precision and accuracy were all below 20% except for 20.27 at the LLOQ of 1 for fentanyl. All other parameters for fentanyl were under 10% well under our 20% cutoff. The ranges for intra-assay variability and error are shown on Table 7 and again fentanyl is the only one with percentages above 20. If only 2 of the 3 days are considered, the percentages drop below 20%, and are even below 15%.

**TABLE 5.** Inter-assay mean and standard deviation (SD) of validation samples.

| Drug / Metabolite | LLOQ (ng/mL) | LQC (ng/mL) | MQC (ng/mL) | HQC (ng/mL) |
| --- | --- | --- | --- | --- |
| | Mean ± SD | Mean ± SD | Mean ± SD | Mean ± SD |
| 6-MAM | 5 ± 0.8 | 14.5 ± 1.9 | 622.3 ± 55.9 | 1928.1 ± 135.3 |
| 7-Amino Clonazepam | 5.6 ± 7.89 | 16.7 ± 1.0 | 598..9 ± 29.3 | 1945.9 ± 161.3 |
| Alprazolam | 5.0 ± 0.6 | 14.6 ± 1.5 | 573.4 ± 32.5 | 19454 ± 148.8 |
| Amitriptyline* | 5.5 ± 0.4 | 16.9 ± 0.8 | 543.9 ± 18.4 | 738.5 ± 40.7 |
| Amphetamine* | 5.4 ± 0.5 | 16.7 ± 0.7 | 581.5 ± 24.6 | 802.4 ± 64.4 |
| a-OH Alprazolam | 4.8 ± 0.6 | 15.5 ± 1.3 | 595.6 ± 28.6 | 1768.6 ± 118.6 |
| Benzoylcegonine | 5.4 ± 0.7 | 16.5 ± 1.1 | 586.7 ± 16.0 | 2069.3 ± 262.1 |
| Buprenorphine | 5.0 ± 0.7 | 15.6 ± 0.9 | 588.1 ± 47.3 | 1814.0 ± 172.7 |
| Carisoprodol | 5.3 ± 0.6 | 16.1 ± 1.4 | 533.7 ± 34.2 | 1709.8 ± 129.1 |
| Chlordiazepoxide* | 5.5 ± 0.7 | 16.5 ± 0.7 | 623.9 ± 34.8 | 778.7 ± 26.3 |
| Clonazepam | 5.2 ± 0.9 | 14.5 ± 1.8 | 585.9 ± 49.5 | 1892.9 ± 63.3 |
| Codeine | 4.8 ± 0.7 | 16.0 ± 1.7 | 597.7 ± 80.4 | 1932.1 ± 333.6 |
| Cotinine | 5.7 ± 0.2 | 15.6 ± 0.5 | 588.5 ± 14.0 | 1883.3 ± 69.5 |
| Cyclobenzaprine | 4.8 ± 0.8 | 15.1 ± 1.4 | 591.8 ± 25.6 | 1839.5 ± 88.0 |
| Desalkylflurazepam | 4.8 ± 0.5 | 15.8 ± 1.7 | 605.7 ± 54.1 | 1954.7 ± 153.3 |
| Desipramine | 5.5 ± 0.3 | 16.0 ± 0.8 | 555.8 ± 38.7 | 1873.5 ± 125.1 |
| Diazepam | 5.2 ± 0.3 | 16.2 ± 1.2 | 603.4 ± 45.0 | 2004.6 ± 231.0 |
| Dihydrocodeine | 5.1 ± 0.8 | 14.5 ± 1.4 | 527.8 ± 43.0 | 1858.0 ± 170.1 |
| Doxepin | 5.1 ± 0.58 | 15.8 ± 0.9 | 587.4 ± 24.3 | 1882.1 ± 74.8 |
| EDDP | 4.8 ± 0.3 | 14.6 ± 1.1 | 520.1 ± 26.4 | 1673.9 ± 127.2 |
| Fentanyl | 0.9 ± 0.4 | 3.0 ± 0.3 | 568.0 ± 28.5 | 1787.3 ± 91.7 |
| Fluoxetine | 5.3 ± 0.6 | 16.2 ± 0.7 | 576.3 ± 34.5 | 1777.0 ± 195.1 |
| Gabapentin | 27.8 ± 3.5 | 75.1 ± 12.3 | 603.0 ± 105.7 | 1767.1 ± 206.4 |
| Hydrocodone | 5.7 ± 0.8 | 15.9 ± 1.8 | 599.2 ± 61.4 | 1753.3 ± 86.4 |
|  | Mean ± SD | Mean ± SD | Mean ± SD | Mean ± SD |
| Hydromorphone | 5.1 ± 0.7 | 14.2 ± 1.6 | 591.5 ± 89.0 | 1873.8 ± 232.1 |
| Imipramine | 4.8 ± 0.7 | 15.0 ± 1.1 | 630.2 ± 38.0 | 2040.0 ± 131.8 |
| Ketamine* | 5.5 ± 0.4 | 16.5 ± 1.0 | 526.2 ± 49.9 | 734.9 ± 78.5 |
| Lorazepam | 5.2 ± 0.8 | 15.8 ± 1.4 | 571.2 ± 25.2 | 1836.7 ± 139.7 |
| MDA* | 5.2 ± 0.6 | 16.1 ± 1.3 | 606.6 ± 39.2 | 830.1 ± 41.4 |
| MDMA | 5.0 ± 0.4 | 14.4 ± 1.1 | 532.9 ± 40.2 | 1693.1 ± 173.0 |
| MDPV | 5.3 ± 0.4 | 15.8 ± 1.2 | 602.1 ± 29.8 | 1894.4 ± 134.5 |
| Meperidine* | 5.3 ± 0.3 | 15.9 ± 0.6 | 579.7 ± 23.8 | 773.3 ± 23.6 |
| Meprobamate | 5.3 ± 0.5 | 15.6 ± 0.7 | 583.1 ± 23.2 | 1889.6 ± 87.3 |
| Methadone | 5.3 ± 0.4 | 17.1 ± 0.5 | 570.2 ± 54.9 | 1685.8 ± 138.8 |
| Methamphetamine | 5.1 ± 0.4 | 15.1 ± 0.5 | 585.6 ± 25.9 | 1837 ± 85.7 |
| Methylphenidate* | 5.6 ± 0.3 | 16.6 ± 0.7 | 586.0 ± 26.1 | 790.3 ± 26.9 |
| Morphine | 5.6 ± 0.6 | 16.6 ± 2.1 | 568.8 ± 67.9 | 1763.8 ± 202.8 |
| Naloxone | 5.2 ± 0.6 | 15.4 ± 2.1 | 584.0 ± 48.1 | 1698.5 ± 109.7 |
| Naltrexone | 5.2 ± 0.7 | 15.7 ± 1.6 | 566.2 ± 34.8 | 1868.2 ± 172.0 |
| Nicotine | 5.0 ± 0.3 | 15.3 ± 0.6 | 592.2 ± 30.0 | 1886.8 ± 150.1 |
| Norbuprenorphine | 4.9 ± 0.5 | 15.0 ± 1.3 | 590.8 ± 20.9 | 1814.0 ± 130.7 |
| Nordiazepam | 4.8 ± 0.5 | 15.5 ± 1.6 | 571.6 ± 43.9 | 1836.4 ± 122.4 |
| Norfentanyl | 5.3 ± 0.5 | 16.4 ± 1.8 | 619.4 ± 50.6 | 1745.5 ± 128.5 |
| Norhydrocodone | 5.1 ± 0.6 | 16.1 ± 2.1 | 635.8 ± 58.4 | 1752.7 ± 138.4 |
| Norketamine | 5.1 ± 0.4 | 14.6 ± 1.4 | 573.3 ± 37.5 | 1732.5 ± 74.5 |
| Noroxycodone | 5.1 ± 0.8 | 16.5 ± 1.4 | 575 ± 76.0 | 1735.3 ± 94.0 |
| Noroxymorphone | 5.4 ± 0.7 | 16.2 ± 1.7 | 644.1 ± 45.0 | 1792.5 ± 92.2 |
| Nortriptyline | 5.1 ± 0.4 | 15.4 ± 1.7 | 521.5 ± 48.6 | 1656.2 ± 255.8 |
| O-desmethyl Tramadol | 5.5 ± 0.4 | 16.2 ± 1.2 | 555.1 ± 33.2 | 1786.9 ± 100.9 |
| Oxazepam | 5.4 ± 0.3 | 17.1 ± 0.6 | 583.6 ± 21.6 | 1895.6 ± 53.4 |
| Oxycodone | 5.0 ± 0.6 | 14.9 ± 1.4 | 579.5 ± 54.3 | 1722.4 ± 82.0 |
| | Mean ± SD | Mean ± SD | Mean ± SD | Mean ± SD |
| Oxymorphone | 5.1 ± 0.7 | 15.0 ± 1.6 | 580.0 ± 37.5 | 1816.0 ± 84.3 |
| Paroxetine | 5.5 ± 0.6 | 16.6 ± 0.9 | 568.4 ± 19.3 | 1842.8160.8 |
| Phencyclidine | 4.8 ± 0.6 | 14.5 ± 1.0 | 574.2 ± 24.9 | 1814.1 ± 77.7 |
| Pregabalin | 26.2 ± 3.9 | 78.5 ± 12.6 | 571.3 ± 78.8 | 1749.2 ± 188.5 |
| Ritalinic Acid* | 5.2 ± 0.9 | 16.0 ± 1.5 | 581.5 ± 58.6 | 782.9 ± 30.5 |
| Tapentadol | 5.6 ± 0.4 | 16.1 ± 1.2 | 580.0 ± 28.6 | 1863.8 ± 60.7 |
| Temazepam | 5.1 ± 0.8 | 17.2 ± 1.8 | 623.5 ± 29.6 | 1813.8 ± 130.5 |
| Tramadol* | 5.1 ± 0.5 | 16.1 ± 0.8 | 572.2 ± 24.3 | 784.7 ± 33.2 |
| Venlafaxine | 5.2 ± 0.6 | 16.0 ± 0.4 | 562.3 ± 41.2 | 1782.0 ± 78.7 |
| Zaleplon | 5.3 ± 0.4 | 16.4 ± 0.9 | 584.1 ± 34.5 | 1749.2 ± 201.4 |
| Zolpidem | 5.0 ± 0.2 | 15.4 ± 0.6 | 574.6 ± 19.6 | 1850.5 ± 91.8 |
| Zopiclone | 5.4 ± 0.4 | 15.7 ± 1.5 | 587.5 ± 57.7 | 1815.9 ± 94.2 |
| THC-COOH | 0.94 ± 0.12 | 2.94 ± 0.30 | 650.9 ± 65.5 | 1828.2 ± 107.2 |
| D-Amphetamine | 2.8 ± 0.1 | 8.3 ± 0.4 | 295.5 ± 14.8 | 940.3 ± 33.4 |
| L-Amphetamine | 2.8 ± 0.1 | 8.4 ± 0.3 | 294.5 ± 6.1 | 874.7 ± 25.7 |
| D-Methamphetamine | 2.6 ± 0.1 | 7.6 ± 0.3 | 300.6 ± 12.1 | 999.5 ± 48.9 |
| L-Methamphetamine | 2.6 ± 0.1 | 7.7 ± 0.4 | 301.6 ± 16.1 | 1013.9 ± 51.4 |

*Note:* Lower limit of quantitation (LLOQ), low quality control (LQC), mid quality control (MQC), high quality control (HQC). Gabapentin and pregabalin had a range from 25 to 2000 ng/mL while the D- and L- assay analytes had a range from 2.5 to 1000 ng/mL

**TABLE 6.** Inter-assay precision and accuracy over 3 days with replicates of 6 for a total of 18 samples

| Drug / Metabolite | LLOQ |  | LQC |  | MQC |  | HQC |  |
| --- | --- | --- | --- | --- | --- | --- | --- | --- |
|  | %CV | %E | %CV | %E | %CV | %E | %CV | %E |
| 6-MAM | 16.74 | -0.81 | 13.23 | -3.33 | 8.99 | 3.72 | 7.01 | 7.12 |
| 7-Amino Clonazepam | 7.89 | 11.06 | 5.85 | 11.07 | 4.88 | -0.18 | 8.29 | 8.10 |
| Alprazolam | 12.90 | -0.43 | 9.98 | -2.78 | 5.67 | -4.43 | 7.65 | 8.05 |
| Amitriptyline* | 6.93 | 10.66 | 4.83 | 12.50 | 3.38 | -9.35 | 5.50 | -7.68 |
| Amphetamine* | 9.25 | 7.37 | 4.31 | 11.13 | 4.22 | -3.09 | 8.02 | 0.30 |
| a-OH Alprazolam | 12.85 | -3.07 | 8.17 | 3.53 | 4.80 | -0.73 | 6.71 | -1.74 |
| Benzoylcegonine | 13.11 | 7.70 | 6.47 | 10.27 | 2.72 | -2.22 | 8.48 | 11.00 |
| Buprenorphine | 14.36 | -0.4 | 5.91 | 4.00 | 8.03 | -1.98 | 9.52 | 0.78 |
| Carisoprodol | 11.91 | 5.21 | 8.89 | 7.49 | 6.41 | -11.05 | 7.55 | -5.01 |
| Chlordiazepoxide* | 7.14 | 9.61 | 4.10 | 9.67 | 5.58 | 3.98 | 3.37 | -2.67 |
| Clonazepam | 16.14 | 0.00 | 12.17 | -3.44 | 8.45 | -2.36 | 3.34 | 5.16 |
| Codeine | 13.99 | -3.39 | 10.35 | 6.82 | 13.45 | -0.38 | 17.27 | 7.34 |
| Cotinine | 4.06 | 5.51 | 3.00 | 3.79 | 2.38 | -1.92 | 3.69 | 4.63 |
| Cyclobenzaprine | 16.84 | -3.26 | 9.20 | 0.56 | 4.33 | -1.36 | 4.78 | 2.19 |
| Desalkylflurazepam | 10.04 | -3.20 | 10.77 | 5.59 | 8.93 | 0.94 | 7.84 | 8.59 |
| Desipramine | 6.20 | 9.72 | 5.00 | 6.49 | 6.95 | -7.37 | 6.68 | 4.08 |
| Diazepam | 5.41 | 4.07 | 7.13 | 8.04 | 7.46 | 0.56 | 11.53 | 11.37 |
| Dihydrocodeine | 11.53 | 11.37 | 9.99 | -3.58 | 8.14 | -12.03 | 9.16 | 3.22 |
| Doxepin | 9.43 | 2.88 | 5.60 | 5.47 | 4.14 | -2.10 | 3.97 | 4.56 |
| EDDP | 6.33 | -3.09 | 7.20 | -2.42 | 5.08 | -13.31 | 7.60 | -7.01 |
| Fentanyl | 20.27 | -5.56 | 8.52 | 0.98 | 5.01 | -5.34 | 5.13 | -0.71 |
| Fluoxetine | 11.39 | 6.41 | 4.06 | 7.76 | 5.99 | -3.95 | 10.98 | -1.28 |
| Gabapentin | 12.44 | 11.22 | 16.40 | 0.18 | 17.52 | 0.50 | 11.68 | -1.83 |
| Hydrocodone | 13.86 | 9.95 | 11.39 | 5.69 | 10.42 | -1.79 | 4.93 | -2.59 |
|  | %CV | %E | %CV | %E | %CV | %E | %CV | %E |
| Hydromorphone | 14.26 | 1.71 | 11.41 | -5.19 | 15.05 | -1.42 | 12.39 | 4.10 |
| Imipramine | 15.02 | -3.06 | 7.36 | 0.10 | 6.02 | 5.04 | 6.46 | 13.33 |
| Ketamine* | 8.04 | 10.31 | 6.06 | 9.84 | 9.49 | -12.30 | 10.68 | -8.14 |
| Lorazepam | 15.07 | 3.43 | 8.96 | 5.01 | 4.41 | -4.81 | 7.55 | 2.04 |
| MDA* | 12.07 | 3.29 | 8.28 | 7.08 | 6.47 | 1.10 | 4.98 | 3.76 |
| MDMA | 8.48 | -0.69 | 7.68 | -4.10 | 7.55 | -11.18 | 10.22 | -5.94 |
| MDPV | 6.69 | 6.29 | 7.75 | 5.64 | 4.94 | 0.34 | 7.10 | 5.24 |
| Meperidine* | 5.76 | 6.24 | 4.06 | 5.95 | 4.10 | -3.38 | 3.05 | -3.34 |
| Meprobamate | 8.88 | 5.12 | 4.37 | 3.87 | 3.98 | -2.81 | 4.62 | 4.98 |
| Methadone | 6.97 | 6.26 | 3.12 | 14.04 | 9.63 | -4.97 | 8.24 | -6.34 |
| Methamphetamine | 8.52 | 2.81 | 3.34 | 0.69 | 4.42 | -2.40 | 4.66 | 2.08 |
| Methylphenidate* | 5.64 | 12.0 | 4.43 | 10.62 | 4.45 | -2.33 | 2.40 | 4.10 |
| Morphine | 12.39 | 4.81 | 12.76 | 10.90 | 11.93 | -5.20 | 11.50 | -2.01 |
| Naloxone | 12.47 | 3.68 | 13.40 | 2.80 | 8.23 | -2.66 | 6.46 | -5.64 |
| Naltrexone | 13.76 | 3.22 | 9.93 | 4.58 | 6.15 | -5.64 | 9.21 | 3.79 |
| Nicotine | 6.21 | -0.71 | 4.17 | 1.96 | 5.06 | -1.31 | 7.95 | 4.82 |
| Norbuprenorphine | 11.01 | -2.41 | 8.94 | -0.29 | 3.54 | -1.53 | 7.20 | 0.78 |
| Nordiazepam | 9.53 | -4.92 | 10.19 | 3.40 | 7.68 | -4.74 | 6.66 | 2.02 |
| Norfentanyl | 10.07 | 5.02 | 10.82 | 9.59 | 8.17 | 3.24 | 7.36 | -3.03 |
| Norhydrocodone | 10.88 | 2.93 | 12.81 | 7.52 | 9.18 | 5.96 | 7.89 | -2.63 |
| Norketamine | 8.65 | 2.39 | 9.55 | -2.93 | 6.54 | -4.45 | 4.30 | -3.75 |
| Noroxycodone | 15.63 | 2.94 | 8.64 | 9.88 | 13.21 | -4.16 | 5.41 | -3.60 |
| Noroxymorphone | 13.80 | 8.37 | 10.32 | 7.91 | 6.99 | 7.35 | 5.15 | -0.42 |
| Nortriptyline | 7.56 | 1.37 | 10.83 | 2.39 | 9.32 | -13.09 | 15.45 | -7.99 |
| O-desmethyl Tramadol | 7.79 | 10.17 | 7.33 | 8.21 | 5.98 | -7.48 | 5.65 | -0.73 |
| Oxazepam | 6.46 | 7.36 | 3.64 | 13.71 | 3.70 | -2.74 | 2.81 | 5.31 |
| Oxycodone | 11.93 | 0.18 | 9.15 | -0.84 | 9.38 | -3.41 | 4.76 | -4.31 |
|  | %CV | %E | %CV | %E | %CV | %E | %CV | %E |
| Oxymorphone | 14.01 | 1.31 | 10.89 | 0.12 | 6.47 | -3.34 | 4.64 | 0.89 |
| Paroxetine | 11.45 | 9.83 | 5.30 | 10.52 | 3.40 | -5.26 | 8.72 | 2.38 |
| Phencyclidine | 11.97 | -3.42 | 6.86 | -3.58 | 4.34 | -4.30 | 4.28 | 0.78 |
| Pregabalin | 14.93 | 4.96 | 16.12 | 4.62 | 13.80 | -4.79 | 10.78 | -2.82 |
| Ritalinic Acid* | 16.35 | 4.00 | 9.65 | 6.73 | 10.07 | -3.08 | 3.89 | -2.13 |
| Tapentadol | 8.40 | 1.72 | 7.34 | 7.14 | 4.93 | -3.34 | 3.26 | 3.55 |
| Temazepam | 15.56 | 1.18 | 10.34 | 14.88 | 4.75 | 3.91 | 7.19 | 0.77 |
| Tramadol | 9.86 | 2.58 | 5.26 | 7.07 | 4.24 | -4.63 | 4.23 | -1.91 |
| Venlafaxine | 11.12 | 3.08 | 2.38 | 6.69 | 7.33 | -6.28 | 4.42 | -1.00 |
| Zaleplon | 6.67 | 6.56 | 5.48 | 9.54 | 5.90 | -2.64 | 11.51 | -2.82 |
| Zolpidem | 4.43 | 0.89 | 3.62 | 2.95 | 3.41 | -4.24 | 4.96 | 2.80 |
| Zopiclone | 6.74 | 8.09 | 9.63 | 4.43 | 9.81 | -2.09 | 5.18 | 0.89 |
| THC-COOH | 12.59 | -5.52 | 10.35 | -2.07 | 10.06 | 8.49 | 5.88 | 1.56 |
| D-Amphetamine | 3.95 | 13.09 | 4.54 | 10.70 | 1.80 | -1.52 | 3.55 | 4.47 |
| L-Amphetamine | 4.12 | 10.00 | 3.04 | 11.52 | 2.07 | -1.85 | 2.94 | -2.81 |
| D-Methamphetamine | 4.44 | 3.17 | 3.98 | 1.30 | 4.04 | 0.21 | 4.89 | 11.05 |
| L-Methamphetamine | 5.54 | 3.21 | 4.98 | 2.33 | 5.32 | 0.52 | 5.07 | 12.66 |

*Note:* lower limit of quantitation (LLOQ), low quality control (LQC), mid quality control (MQC), high quality control (HQC), percent coefficient of variability(%CV), percent error (%E)

**TABLE 7.** Intra-assay Precision and Accuracy: Precision and Accuracy over 3 days with replicates of 6 for each day

| Drug / Metabolite | LLOQ |  |  |  | LQC |  |  |  | MQC |  |  |  | HQC |  |  |  |
| --- | --- | --- | --- | --- | --- | --- | --- | --- | --- | --- | --- | --- | --- | --- | --- | --- |
|  | %CV |  | %E |  | %CV |  | %E |  | %CV |  | %E |  | %CV |  | %E |  |
|  | MIN | MAX | MIN | MAX | MIN | MAX | MIN | MAX | MIN | MAX | MIN | MAX | MIN | MAX | MIN | MAX |
| 6-MAM | 9.01 | 18.07 | -15.64 | 12.17 | 7.28 | 13.63 | -12.10 | 4.12 | 4.86 | 9.74 | -3.09 | 11.41 | 2.58 | 4.71 | -0.05 | 15.12 |
| 7-Amino Clonazepam | 4.33 | 9.39 | 6.00 | 14.87 | 5.44 | 6.65 | 8.68 | 12.41 | 3.32 | 6.00 | -3.03 | 2.52 | 3.27 | 6.43 | -1.10 | 15.50 |
| Alprazolam | 8.16 | 13.14 | -11.60 | 7.20 | 3.35 | 8.88 | -8.82 | 7.32 | 3.12 | 6.64 | -6.64 | -1.90 | 4.77 | 6.81 | 1.42 | 14.77 |
| Amitriptyline* | 3.51 | 11.52 | 9.13 | 11.90 | 4.30 | 5.39 | 9.67 | 14.33 | 2.06 | 3.58 | -11.29 | -7.38 | 2.46 | 6.28 | -10.36 | -3.09 |
| Amphetamine* | 2.28 | 10.19 | -2.20 | 13.20 | 2.72 | 5.62 | 9.07 | 12.43 | 2.06 | 5.57 | -5.07 | -0.91 | 3.46 | 8.89 | -5.00 | 6.59 |
| a-OH Alprazolam | 8.82 | 16.64 | -5.43 | -1.50 | 6.06 | 9.67 | -1.69 | 6.63 | 3.30 | 5.94 | -1.80 | 0.36 | 3.15 | 7.31 | -6.77 | 1.76 |
| Benzoylcegonine | 3.95 | 10.99 | -3.30 | 17.00 | 4.07 | 8.20 | 6.22 | 15.23 | 2.13 | 3.39 | -3.23 | -1.16 | 2.48 | 5.94 | 3.38 | 23.18 |
| Buprenorphine | 8.06 | 17.33 | -7.13 | 10.33 | 2.54 | 8.69 | 2.84 | 6.02 | 3.76 | 6.68 | -10.06 | 3.63 | 2.14 | 9.87 | -7.80 | 8.99 |
| Carisoprodol | 6.63 | 10.14 | -2.07 | 17.70 | 6.03 | 9.21 | 1.10 | 11.82 | 3.99 | 4.47 | -16.33 | -5.62 | 3.79 | 5.61 | -11.33 | 1.84 |
| Chlordiazepoxide* | 3.55 | 10.32 | 8.03 | 10.73 | 2.62 | 3.99 | 5.66 | 11.91 | 2.62 | 3.58 | -2.90 | 7.70 | 1.55 | 3.97 | -4.65 | -0.60 |
| Clonazepam | 2.17 | 7.66 | -17.17 | 19.65 | 4.59 | 10.69 | -12.53 | 8.71 | 2.52 | 7.67 | -8.92 | 6.07 | 1.01 | 3.80 | 1.65 | 7.30 |
| Codeine | 5.91 | 14.73 | -12.08 | 3.07 | 6.84 | 9.35 | -3.67 | 13.09 | 7.06 | 10.54 | -14.33 | 10.37 | 2.72 | 13.64 | -8.97 | 18.21 |
| Cotinine | 1.06 | 3.97 | 0.77 | 9.03 | 0.85 | 2.21 | 0.33 | 6.51 | 1.79 | 3.14 | -2.90 | -0.74 | 2.57 | 3.04 | 1.37 | 7.69 |
| Cyclobenzaprine | 10.47 | 19.32 | -15.93 | 5.40 | 5.85 | 8.78 | -5.98 | 6.97 | 2.48 | 3.82 | -4.49 | 2.69 | 2.10 | 3.45 | -3.46 | 6.03 |
| Desalkylflurazepam | 5.97 | 8.27 | -8.63 | 6.70 | 5.69 | 13.22 | -3.92 | 12.83 | 4.34 | 6.04 | -8.98 | 7.65 | 2.60 | 5.00 | -1.48 | 15.23 |
| Desipramine | 5.16 | 7.14 | 8.00 | 11.50 | 2.18 | 7.35 | 4.69 | 9.53 | 1.28 | 7.21 | -13.39 | -1.22 | 2.75 | 3.75 | -1.82 | 9.98 |
| Diazepam | 3.25 | 5.61 | -0.30 | 6.30 | 3.58 | 8.12 | 1.24 | 13.38 | 2.11 | 4.41 | -5.58 | 9.68 | 5.38 | 11.58 | 3.58 | 23.59 |
| Dihydrocodeine | 10.04 | 16.88 | -9.40 | 12.63 | 7.98 | 13.09 | -7.00 | -1.78 | 4.84 | 10.52 | -13.49 | -10.99 | 4.55 | 9.20 | -3.55 | 9.85 |
| Doxepin | 7.00 | 12.15 | 0.10 | 7.27 | 4.92 | 5.47 | 2.13 | 8.61 | 2.31 | 4.47 | -4.49 | 0.38 | 2.80 | 6.07 | 4.07 | 5.02 |
| EDDP | 5.06 | 6.14 | -7.17 | 1.37 | 4.29 | 9.80 | -5.34 | 1.72 | 2.75 | 3.37 | -16.90 | -8.59 | 3.09 | 4.92 | -14.97 | -0.45 |
| Fentanyl | 14.53 | 23.15 | -26.75 | 1.67 | 7.03 | 8.83 | -4.89 | 4.33 | 1.46 | 3.85 | -8.80 | -0.24 | 3.00 | 4.70 | -5.22 | 2.49 |
| Fluoxetine | 5.24 | 14.21 | 3.50 | 8.33 | 2.79 | 5.22 | 6.56 | 8.46 | 3.20 | 5.05 | -8.42 | 1.90 | 5.71 | 7.03 | -9.96 | 10.59 |
| Gabapentin | 4.76 | 17.33 | 4.87 | 19.29 | 6.99 | 13.05 | -15.95 | 13.29 | 6.74 | 12.49 | -16.17 | 19.80 | 6.23 | 11.44 | -11.32 | 6.27 |
|  | %CV |  | %E |  | %CV |  | %E |  | %CV |  | %E |  | %CV |  | %E |  |
|  | MIN | MAX | MIN | MAX | MIN | MAX | MIN | MAX | MIN | MAX | MIN | MAX | MIN | MAX | MIN | MAX |
| Hydrocodone | 10.55 | 17.43 | 5.43 | 14.00 | 5.47 | 11.96 | -1.69 | 17.42 | 4.70 | 5.19 | -13.60 | 7.84 | 1.50 | 6.96 | -4.02 | 0.16 |
| Hydromorphone | 8.91 | 16.26 | -5.57 | 12.17 | 10.45 | 12.60 | -6.96 | -2.01 | 4.86 | 18.92 | -11.73 | 11.41 | 2.58 | 13.75 | -2.49 | 15.12 |
| Imipramine | 9.76 | 16.92 | -7.60 | 4.20 | 2.18 | 12.02 | -0.97 | 0.73 | 1.41 | 4.14 | -0.82 | 11.93 | 2.37 | 4.76 | 5.43 | 18.80 |
| Ketamine* | 4.93 | 7.35 | 1.93 | 15.50 | 6.04 | 6.77 | 8.97 | 10.61 | 3.43 | 4.18 | -18.13 | -1.77 | 3.65 | 5.77 | -19.13 | 1.71 |
| Lorazepam | 1.11 | 16.90 | -7.33 | 13.15 | 4.27 | 11.16 | 0.73 | 8.02 | 3.08 | 3.64 | -8.77 | -2.71 | 3.59 | 7.28 | -4.13 | 8.19 |
| MDA* | 9.43 | 11.52 | -4.93 | 13.07 | 5.30 | 11.03 | 4.78 | 10.87 | 3.40 | 9.79 | 0.09 | 2.31 | 2.21 | 4.70 | 1.37 | 8.37 |
| MDMA | 5.74 | 8.93 | -7.23 | 4.70 | 3.36 | 8.15 | -9.64 | 2.56 | 3.15 | 5.42 | -17.29 | -4.25 | 3.84 | 6.78 | -12.90 | 5.72 |
| MDPV | 3.00 | 11.27 | 4.93 | 8.77 | 4.60 | 6.19 | -2.82 | 10.82 | 4.20 | 4.86 | -2.45 | 3.66 | 2.27 | 6.38 | -0.25 | 13.82 |
| Meperidine* | 1.20 | 5.32 | -0.20 | 9.53 | 1.82 | 3.58 | 1.72 | 9.02 | 1.48 | 5.68 | -6.50 | -1.04 | 1.21 | 4.51 | -5.16 | -2.42 |
| Meprobamate | 3.72 | 10.44 | 1.83 | 10.83 | 2.06 | 6.39 | 1.39 | 5.50 | 2.61 | 4.33 | -5.00 | -1.06 | 2.69 | 4.46 | 2.27 | 9.30 |
| Methadone | 5.26 | 8.68 | 2.30 | 8.97 | 1.50 | 3.98 | 11.96 | 16.42 | 2.61 | 9.74 | -13.82 | 2.85 | 3.92 | 4.42 | -12.79 | 0.10 |
| Methamphetamine | 4.84 | 9.89 | -2.60 | 7.53 | 1.71 | 4.41 | -0.02 | 1.66 | 1.71 | 2.92 | -6.31 | 2.48 | 2.10 | 4.41 | -2.22 | 6.50 |
| Methylphenidate* | 5.07 | 6.05 | 9.07 | 13.60 | 2.18 | 4.86 | 7.54 | 14.90 | 1.70 | 2.49 | -7.45 | 1.52 | 2.30 | 4.58 | -2.83 | 0.27 |
| Morphine | 9.36 | 13.19 | 0.30 | 13.63 | 8.05 | 10.54 | -1.54 | 21.87 | 6.05 | 8.83 | -17.22 | 4.53 | 7.02 | 9.10 | -10.99 | 9.15 |
| Naloxone | 6.04 | 13.40 | -1.03 | 12.70 | 10.16 | 14.14 | -4.99 | 12.33 | 6.59 | 8.14 | -7.62 | 1.54 | 2.47 | 8.16 | -8.08 | -3.20 |
| Naltrexone | 9.68 | 15.15 | -3.23 | 11.84 | 8.31 | 11.20 | -0.04 | 8.54 | 4.11 | 6.59 | -7.70 | -1.60 | 4.79 | 11.07 | -0.44 | 8.19 |
| Nicotine | 2.98 | 6.09 | -4.40 | 5.10 | 1.49 | 3.80 | -0.34 | 6.23 | 2.27 | 4.29 | -5.74 | 3.74 | 1.67 | 4.22 | -3.38 | 14.85 |
| Norbuprenorphine | 6.96 | 10.42 | -11.27 | 5.90 | 5.11 | 8.76 | -4.94 | 7.59 | 2.29 | 4.50 | -2.15 | -0.41 | 3.79 | 10.56 | -2.01 | 3.85 |
| Nordiazepam | 5.19 | 9.95 | -11.03 | -1.33 | 4.88 | 14.06 | -2.36 | 9.71 | 5.30 | 6.07 | -10.87 | 1.56 | 3.79 | 5.86 | -4.57 | 5.71 |
| Norfentanyl | 7.81 | 11.01 | 0.43 | 10.50 | 8.97 | 11.84 | 6.40 | 15.21 | 6.26 | 9.54 | -1.92 | 6.44 | 5.62 | 8.23 | -7.19 | -0.09 |
| Norhydrocodone | 5.68 | 15.60 | 0.40 | 6.60 | 9.43 | 13.18 | -0.61 | 13.97 | 7.21 | 11.23 | 1.61 | 9.54 | 6.06 | 7.61 | -8.22 | 0.31 |
| Norketamine | 6.55 | 8.72 | -3.83 | 6.40 | 6.17 | 9.01 | -9.44 | 5.31 | 3.55 | 4.31 | -9.08 | 2.41 | 2.00 | 4.92 | -6.43 | -0.59 |
| Noroxycodone | 14.45 | 16.73 | -2.60 | 7.83 | 4.43 | 13.35 | 8.29 | 12.28 | 5.20 | 16.90 | -13.54 | 5.18 | 2.07 | 6.97 | -7.78 | -1.00 |
| Noroxymorphone | 9.38 | 17.85 | 3.73 | 16.33 | 7.28 | 12.74 | 4.22 | 12.24 | 2.88 | 3.33 | 3.52 | 11.98 | 2.55 | 7.08 | -3.14 | 3.01 |
| Nortriptyline | 7.12 | 7.15 | -1.83 | 4.57 | 3.51 | 5.52 | -11.40 | 11.44 | 2.75 | 2.84 | -19.93 | -2.58 | 2.62 | 4.18 | -18.75 | 11.08 |
| O-desmethyl Tramadol | 3.44 | 9.99 | 6.50 | 13.87 | 2.60 | 9.77 | 3.96 | 12.03 | 5.08 | 6.01 | -10.35 | -3.76 | 5.12 | 6.36 | -1.97 | 0.52 |
|  | %CV |  | %E |  | %CV |  | %E |  | %CV |  | %E |  | %CV |  | %E |  |
|  | MIN | MAX | MIN | MAX | MIN | MAX | MIN | MAX | MIN | MAX | MIN | MAX | MIN | MAX | MIN | MAX |
| Oxazepam | 4.77 | 7.79 | 5.20 | 8.43 | 2.35 | 3.72 | 10.57 | 16.98 | 2.30 | 5.13 | -3.76 | -1.67 | 2.31 | 2.45 | 3.73 | 7.78 |
| Oxycodone | 8.27 | 9.76 | -11.23 | 6.23 | 6.54 | 11.45 | -5.59 | 3.09 | 6.01 | 10.16 | -10.54 | 0.54 | 3.27 | 4.99 | -7.15 | -1.19 |
| Oxymorphone | 9.91 | 16.05 | -4.40 | 8.20 | 7.73 | 14.01 | -5.56 | 7.19 | 4.31 | 7.44 | -6.06 | -1.90 | 3.63 | 5.86 | -0.68 | 2.38 |
| Paroxetine | 7.68 | 16.21 | 8.97 | 11.40 | 2.16 | 6.44 | 8.13 | 12.07 | 2.93 | 3.31 | -7.27 | -3.22 | 4.36 | 10.92 | -3.71 | 7.83 |
| Phencyclidine | 6.58 | 15.72 | -9.23 | 4.50 | 4.55 | 7.46 | -8.27 | -1.21 | 2.84 | 3.94 | -8.09 | -1.38 | 3.25 | 4.47 | -0.81 | 3.88 |
| Pregabalin | 11.84 | 17.10 | -1.53 | 10.89 | 8.00 | 17.30 | -7.30 | 14.20 | 9.76 | 17.69 | -10.64 | 0.72 | 5.82 | 15.67 | -7.15 | -0.35 |
| Ritalinic Acid* | 2.91 | 16.92 | -7.03 | 22.63 | 4.01 | 12.30 | -0.56 | 12.41 | 4.22 | 14.76 | -8.74 | 1.37 | 3.68 | 4.52 | -3.08 | -1.18 |
| Tapentadol | 1.94 | 10.80 | -1.73 | 7.03 | 3.82 | 5.42 | -1.72 | 12.80 | 1.62 | 5.14 | -7.71 | -0.41 | 2.72 | 3.87 | 2.32 | 5.17 |
| Temazepam | 5.27 | 12.57 | -15.80 | 9.80 | 4.85 | 7.18 | 6.83 | 27.88 | 1.15 | 2.54 | -2.14 | 7.89 | 1.32 | 4.02 | -7.81 | 7.95 |
| Tramadol | 3.94 | 8.92 | -5.03 | 11.77 | 4.73 | 5.09 | 4.71 | 10.89 | 2.47 | 4.92 | -6.52 | -3.37 | 3.58 | 5.70 | -3.08 | -1.10 |
| Venlafaxine | 5.86 | 7.08 | -10.10 | 10.27 | 1.70 | 2.82 | 5.80 | 8.38 | 3.01 | 5.73 | -14.40 | -1.74 | 1.86 | 2.84 | -5.99 | -0.86 |
| Zaleplon | 4.79 | 6.74 | 1.67 | 9.77 | 0.91 | 6.98 | 5.97 | 13.88 | 4.74 | 6.08 | -5.49 | 1.63 | 4.94 | 14.24 | -13.41 | 5.24 |
| Zolpidem | 2.24 | 5.12 | -1.43 | 4.43 | 2.48 | 3.48 | 0.82 | 6.28 | 1.08 | 2.64 | -7.17 | -0.79 | 1.26 | 3.41 | -1.48 | 7.09 |
| Zopiclone | 4.44 | 9.90 | 6.57 | 10.10 | 7.13 | 8.32 | -3.02 | 12.93 | 3.98 | 7.60 | -13.09 | 6.17 | 2.66 | 3.99 | -3.07 | 4.84 |
| THC-COOH | 5.20 | 7.38 | -17.50 | 5.17 | 3.82 | 10.03 | -13.78 | 4.94 | 2.67 | 6.62 | -6.60 | 20.04 | 1.18 | 8.33 | -4.49 | 6.79 |
| D-Amphetamine | 1.38 | 3.51 | 8.44 | 15.79 | 1.14 | 1.40 | 4.38 | 15.69 | 0.69 | 1.73 | -2.89 | -0.02 | 1.46 | 2.99 | 0.75 | 7.90 |
| L-Amphetamine | 2.60 | 4.47 | 6.63 | 13.33 | 1.11 | 1.79 | 9.38 | 15.71 | 1.14 | 1.57 | -3.13 | 0.31 | 1.46 | 2.33 | -5.16 | 0.02 |
| D-Methamphetamine | 1.28 | 4.42 | -1.56 | 5.73 | 1.67 | 2.66 | -3.19 | 4.96 | 1.75 | 2.21 | -4.28 | 4.24 | 1.67 | 2.74 | 7.59 | 17.89 |
| L-Methamphetamine | 1.66 | 5.19 | -2.83 | 7.65 | 2.59 | 3.56 | -3.14 | 6.69 | 0.55 | 0.99 | -4.74 | 7.49 | 2.17 | 5.87 | 8.68 | 17.91 |

Lower limit of Quantitation (LLOQ), Low Quality Control (LQC), Mid Quality Control (MQC), High Quality Control (HQC), Percent Coefficient of Variability(%CV), Percent error (%E)

### 3.3 Partial volumes accuracy and precision

An MPA surrogate sample was prepared at 4000 ng/mL. To determine the concentration of this sample, a dilution must be made so the final concentration would be less than 2000 ng/mL to get it in the measurement range of the assay. Three replicates of four dilutions were made and tested: 1) 1:5 target 800 ng/mL; 2) 1:10 with a target of 400 ng/mL; 3) 1:20 with a target of 200 ng/mL; and 4) 1:50 with a target of 80 ng/mL. The results shown in Table 8 indicate that a 1:10 dilution is safe for all analytes. Most of the analytes can be diluted at all levels. Notable exceptions are 7-aminoclonazepam and temazepam which only accommodated the 1:10 dilution.

**TABLE 8.**
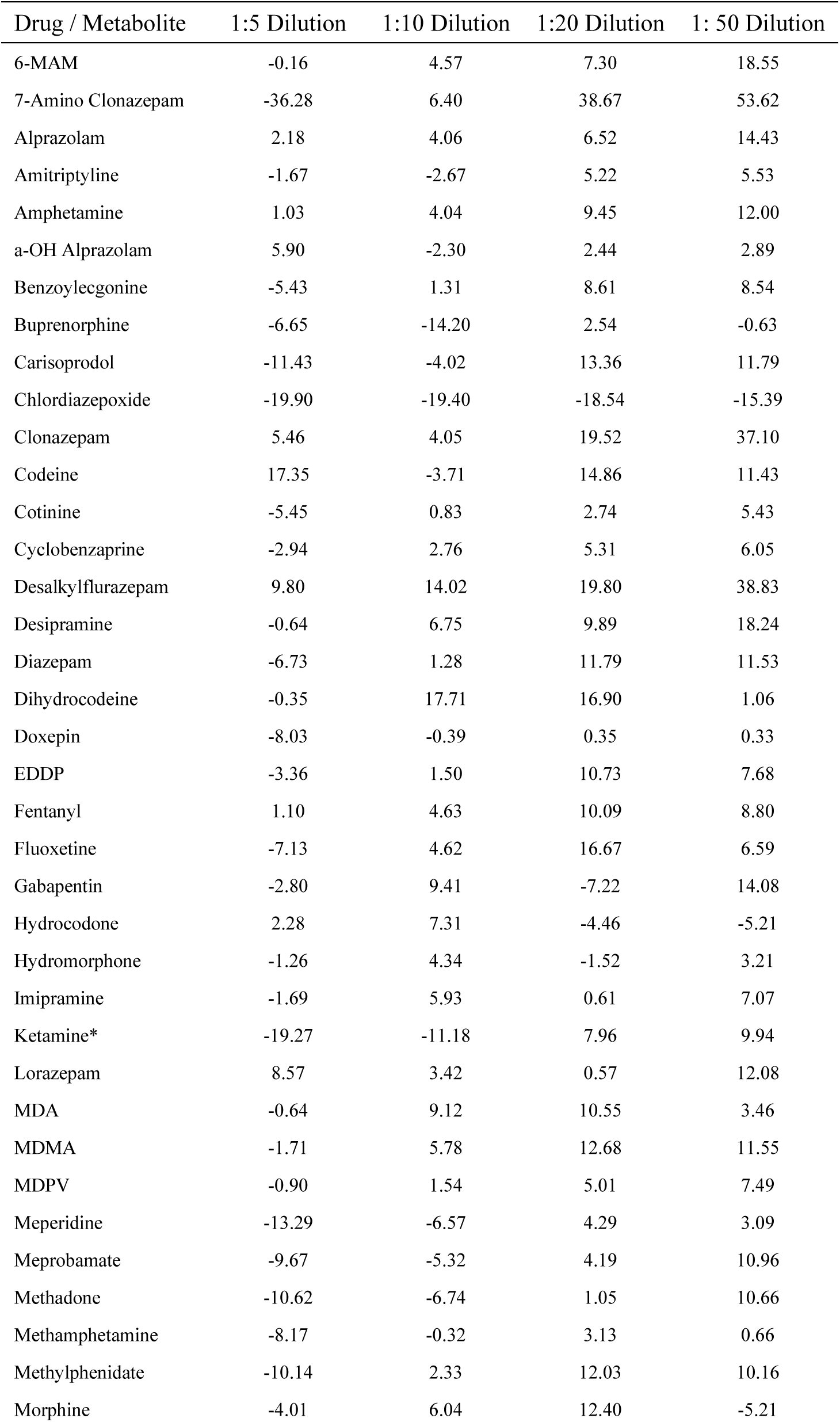
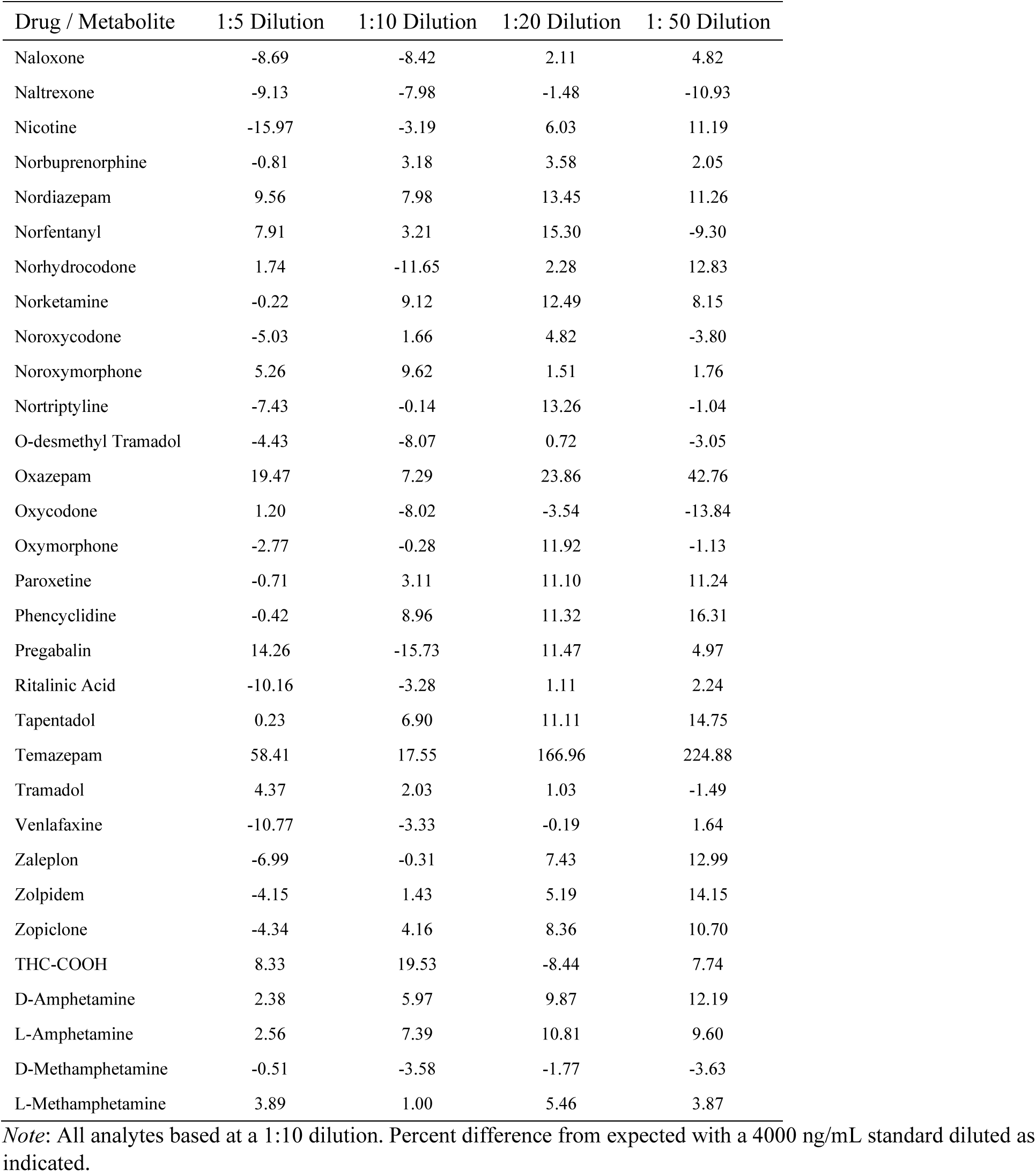
Dilution study

| Drug / Metabolite | 1:5 Dilution | 1:10 Dilution | 1:20 Dilution | 1: 50 Dilution |
| --- | --- | --- | --- | --- |
| 6-MAM | -0.16 | 4.57 | 7.30 | 18.55 |
| 7-Amino Clonazepam | -36.28 | 6.40 | 38.67 | 53.62 |
| Alprazolam | 2.18 | 4.06 | 6.52 | 14.43 |
| Amitriptyline | -1.67 | -2.67 | 5.22 | 5.53 |
| Amphetamine | 1.03 | 4.04 | 9.45 | 12.00 |
| a-OH Alprazolam | 5.90 | -2.30 | 2.44 | 2.89 |
| Benzoylcegonine | -5.43 | 1.31 | 8.61 | 8.54 |
| Buprenorphine | -6.65 | -14.20 | 2.54 | -0.63 |
| Carisoprodol | -11.43 | -4.02 | 13.36 | 11.79 |
| Chlordiazepoxide | -19.90 | -19.40 | -18.54 | -15.39 |
| Clonazepam | 5.46 | 4.05 | 19.52 | 37.10 |
| Codeine | 17.35 | -3.71 | 14.86 | 11.43 |
| Cotinine | -5.45 | 0.83 | 2.74 | 5.43 |
| Cyclobenzaprine | -2.94 | 2.76 | 5.31 | 6.05 |
| Desalkylflurazepam | 9.80 | 14.02 | 19.80 | 38.83 |
| Desipramine | -0.64 | 6.75 | 9.89 | 18.24 |
| Diazepam | -6.73 | 1.28 | 11.79 | 11.53 |
| Dihydrocodeine | -0.35 | 17.71 | 16.90 | 1.06 |
| Doxepin | -8.03 | -0.39 | 0.35 | 0.33 |
| EDDP | -3.36 | 1.50 | 10.73 | 7.68 |
| Fentanyl | 1.10 | 4.63 | 10.09 | 8.80 |
| Fluoxetine | -7.13 | 4.62 | 16.67 | 6.59 |
| Gabapentin | -2.80 | 9.41 | -7.22 | 14.08 |
| Hydrocodone | 2.28 | 7.31 | -4.46 | -5.21 |
| Hydromorphone | -1.26 | 4.34 | -1.52 | 3.21 |
| Imipramine | -1.69 | 5.93 | 0.61 | 7.07 |
| Ketamine* | -19.27 | -11.18 | 7.96 | 9.94 |
| Lorazepam | 8.57 | 3.42 | 0.57 | 12.08 |
| MDA | -0.64 | 9.12 | 10.55 | 3.46 |
| MDMA | -1.71 | 5.78 | 12.68 | 11.55 |
| MDPV | -0.90 | 1.54 | 5.01 | 7.49 |
| Meperidine | -13.29 | -6.57 | 4.29 | 3.09 |
| Meprobamate | -9.67 | -5.32 | 4.19 | 10.96 |
| Methadone | -10.62 | -6.74 | 1.05 | 10.66 |
| Methamphetamine | -8.17 | -0.32 | 3.13 | 0.66 |
| Methylphenidate | -10.14 | 2.33 | 12.03 | 10.16 |
| Morphine | -4.01 | 6.04 | 12.40 | -5.21 |
| Naloxone | -8.69 | -8.42 | 2.11 | 4.82 |
| Naltrexone | -9.13 | -7.98 | -1.48 | -10.93 |
| Nicotine | -15.97 | -3.19 | 6.03 | 11.19 |
| Norbuprenorphine | -0.81 | 3.18 | 3.58 | 2.05 |
| Nordiazepam | 9.56 | 7.98 | 13.45 | 11.26 |
| Norfentanyl | 7.91 | 3.21 | 15.30 | -9.30 |
| Norhydrocodone | 1.74 | -11.65 | 2.28 | 12.83 |
| Norketamine | -0.22 | 9.12 | 12.49 | 8.15 |
| Noroxycodone | -5.03 | 1.66 | 4.82 | -3.80 |
| Noroxymorphone | 5.26 | 9.62 | 1.51 | 1.76 |
| Nortriptyline | -7.43 | -0.14 | 13.26 | -1.04 |
| O-desmethyl Tramadol | -4.43 | -8.07 | 0.72 | -3.05 |
| Oxazepam | 19.47 | 7.29 | 23.86 | 42.76 |
| Oxycodone | 1.20 | -8.02 | -3.54 | -13.84 |
| Oxymorphone | -2.77 | -0.28 | 11.92 | -1.13 |
| Paroxetine | -0.71 | 3.11 | 11.10 | 11.24 |
| Phencyclidine | -0.42 | 8.96 | 11.32 | 16.31 |
| Pregabalin | 14.26 | -15.73 | 11.47 | 4.97 |
| Ritalinic Acid | -10.16 | -3.28 | 1.11 | 2.24 |
| Tapentadol | 0.23 | 6.90 | 11.11 | 14.75 |
| Temazepam | 58.41 | 17.55 | 166.96 | 224.88 |
| Tramadol | 4.37 | 2.03 | 1.03 | -1.49 |
| Venlafaxine | -10.77 | -3.33 | -0.19 | 1.64 |
| Zaleplon | -6.99 | -0.31 | 7.43 | 12.99 |
| Zolpidem | -4.15 | 1.43 | 5.19 | 14.15 |
| Zopiclone | -4.34 | 4.16 | 8.36 | 10.70 |
| THC-COOH | 8.33 | 19.53 | -8.44 | 7.74 |
| D-Amphetamine | 2.38 | 5.97 | 9.87 | 12.19 |
| L-Amphetamine | 2.56 | 7.39 | 10.81 | 9.60 |
| D-Methamphetamine | -0.51 | -3.58 | -1.77 | -3.63 |
| L-Methamphetamine | 3.89 | 1.00 | 5.46 | 3.87 |

*Note:* All analytes based at a 1:10 dilution. Percent difference from expected with a 4000 ng/mL standard diluted as indicated.

### 3.4 Analyte stability and other characteristics

QC samples were subjected to several conditions to test the stability of the analytes as shown in Table 9.

**TABLE 9.**
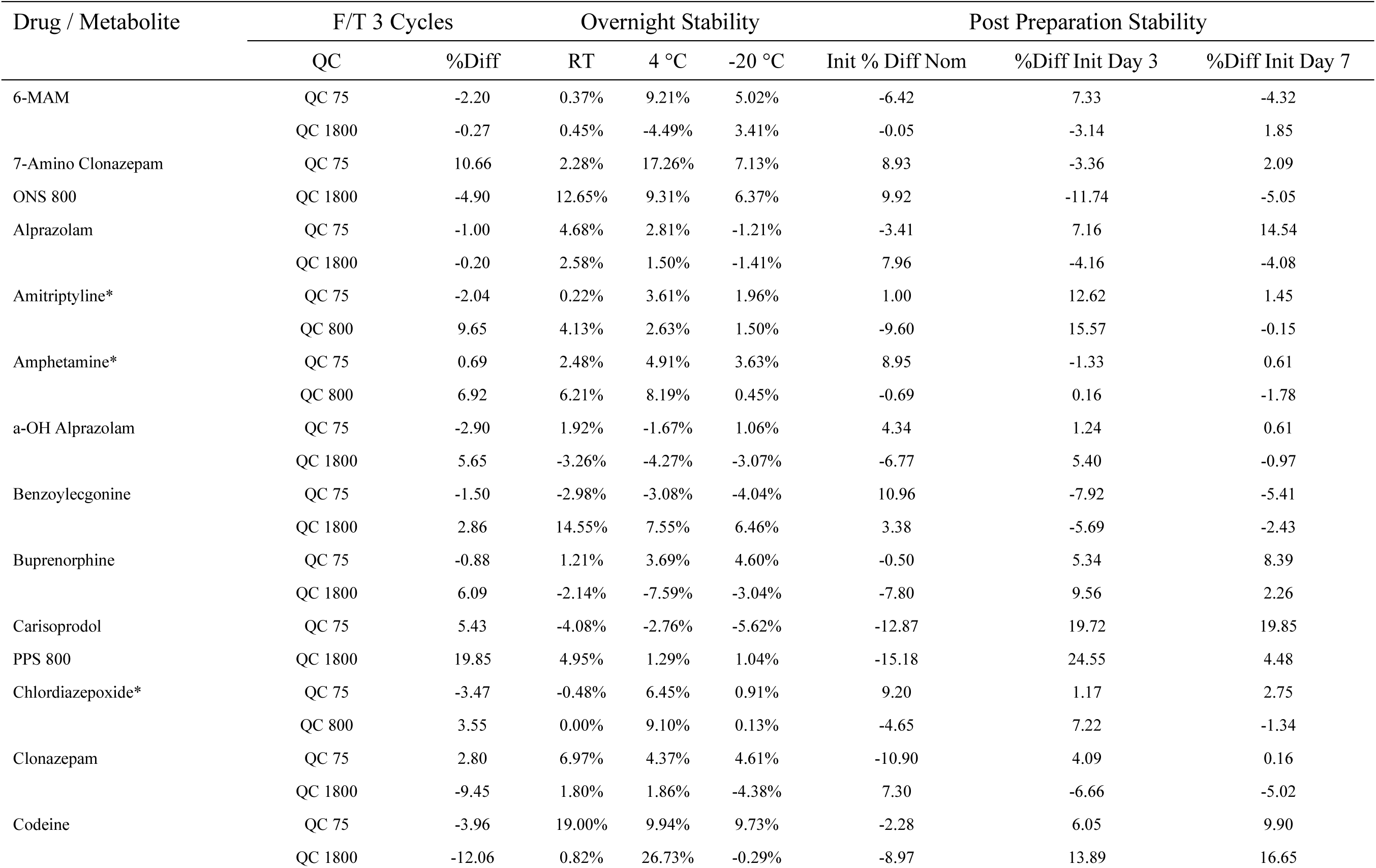
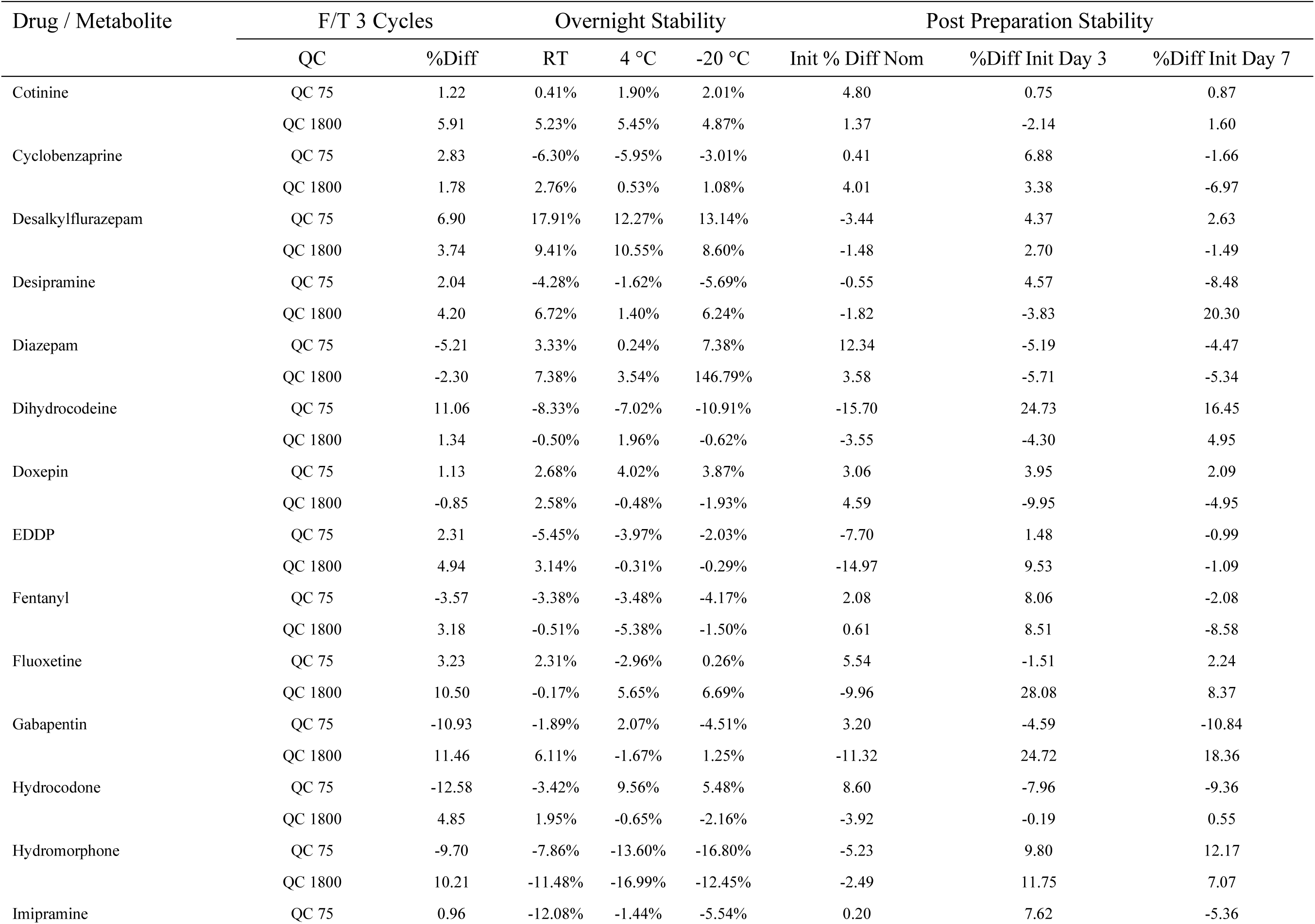
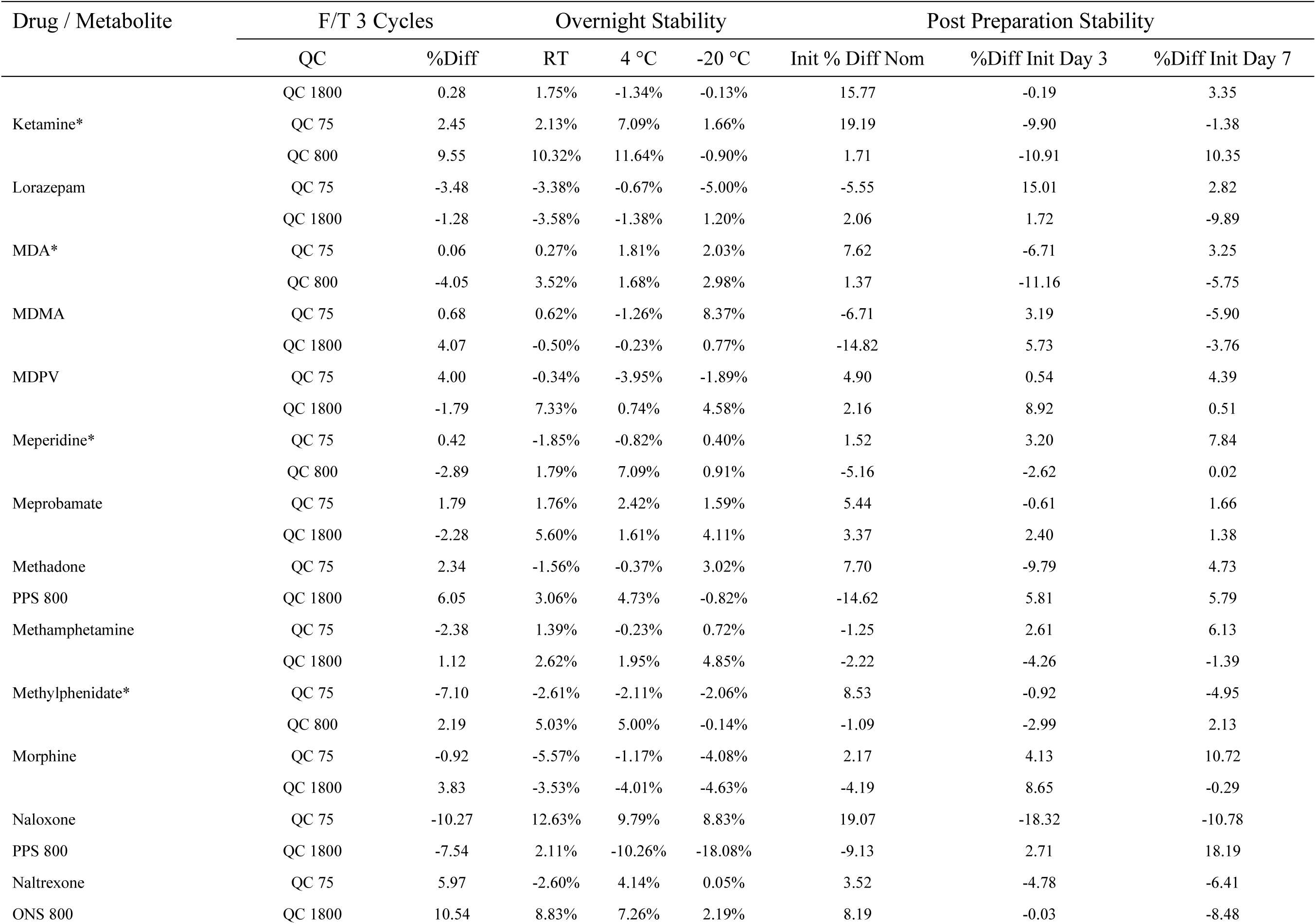
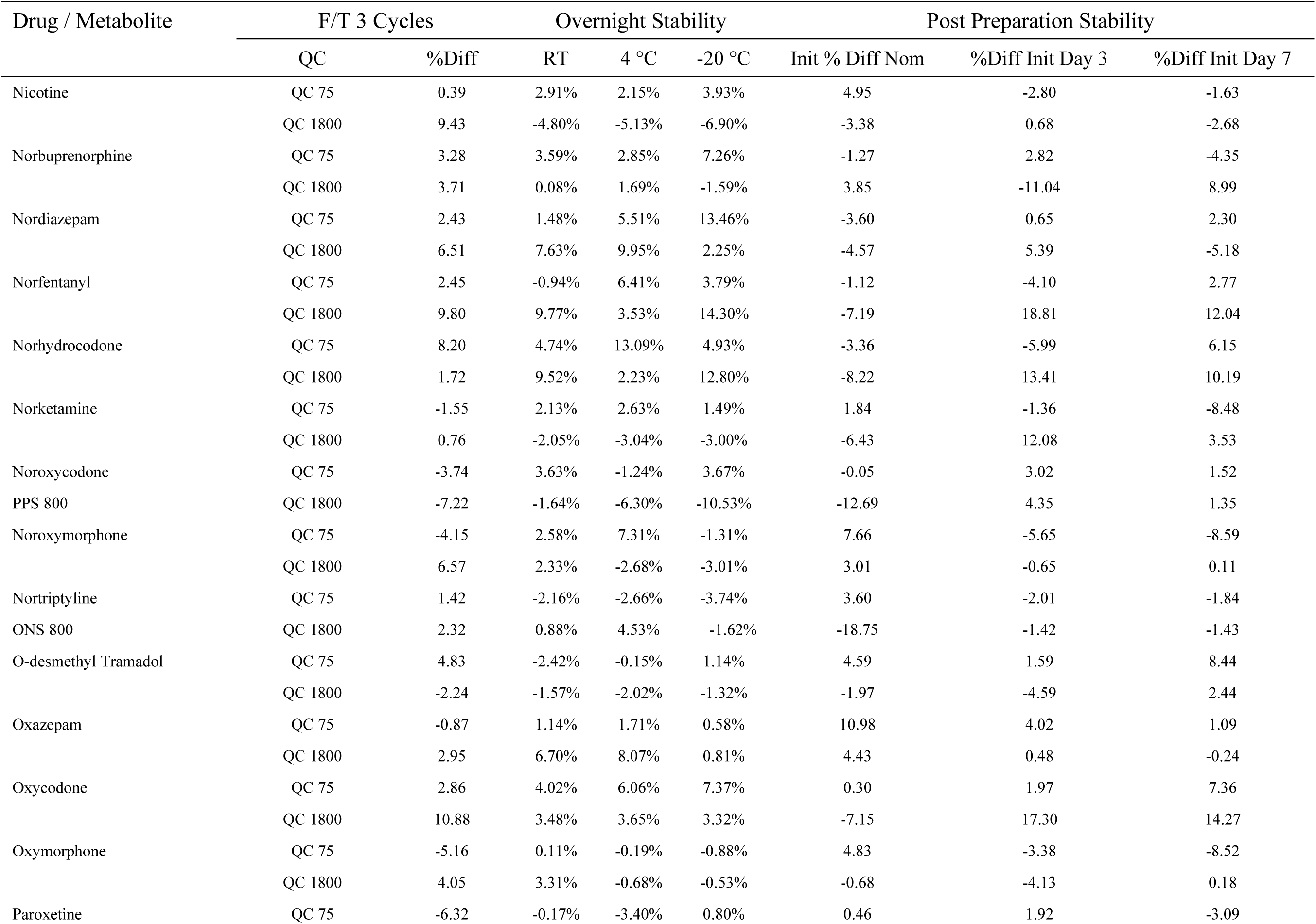
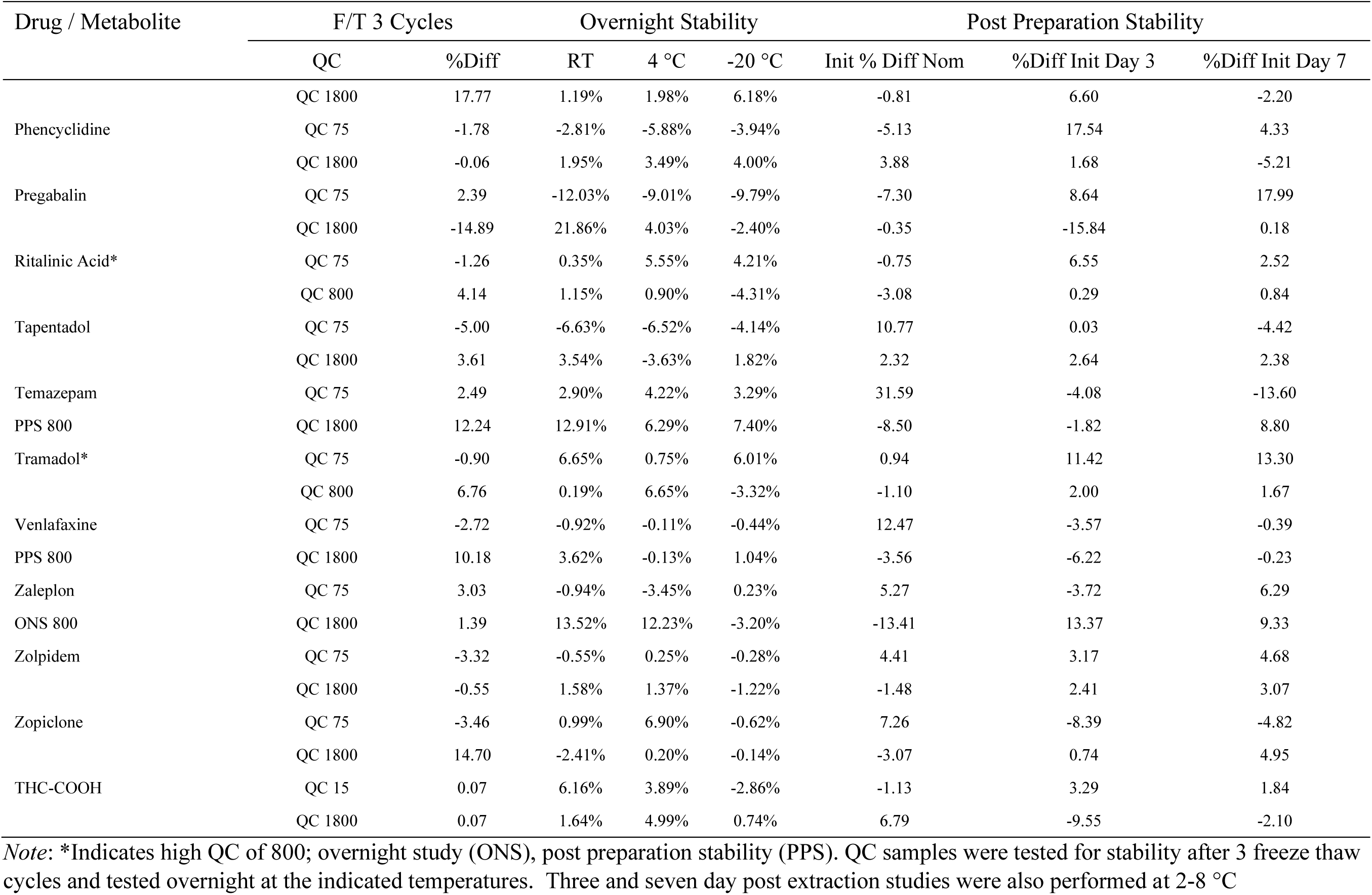
Stability testing

**TABLE 10.**
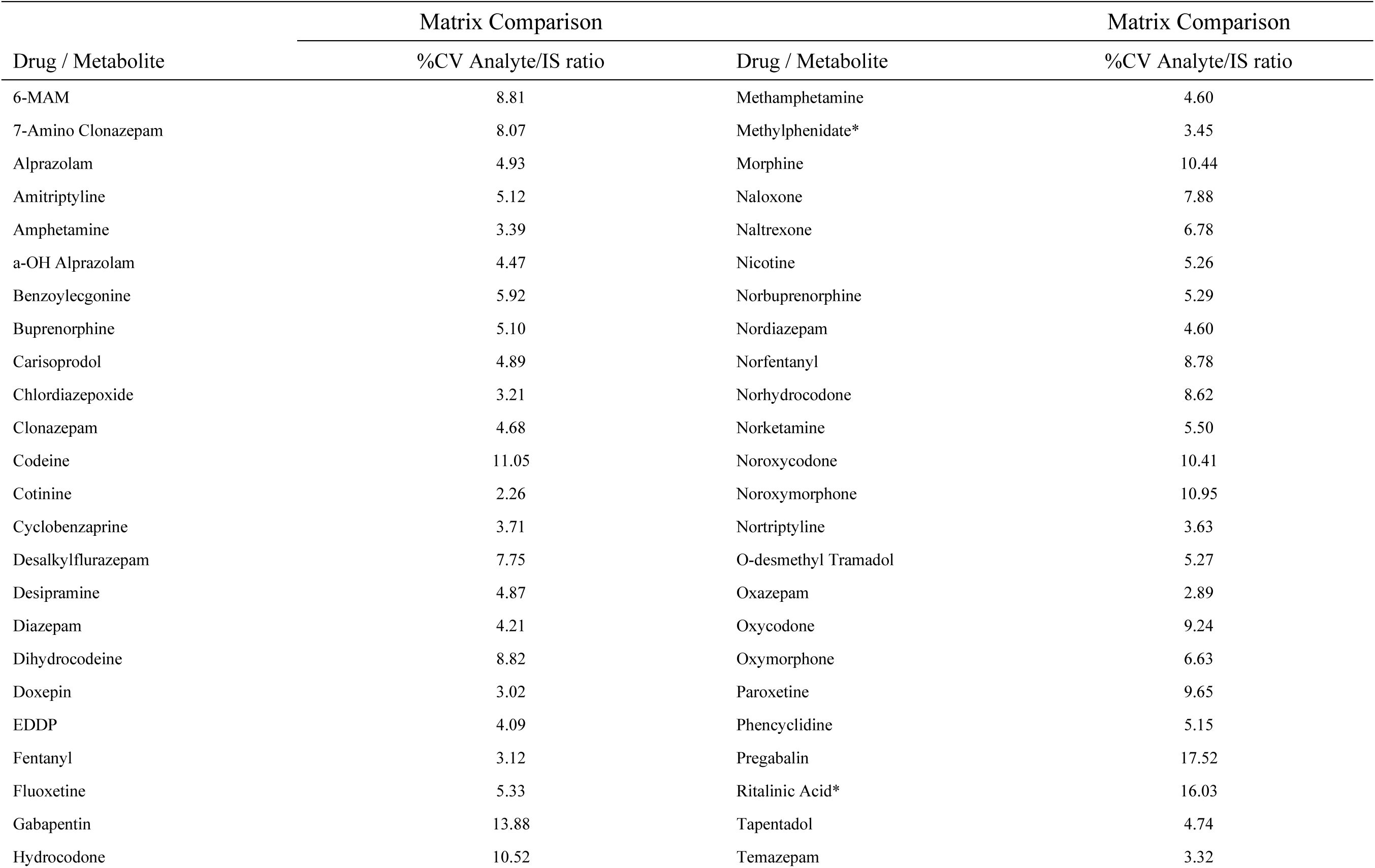
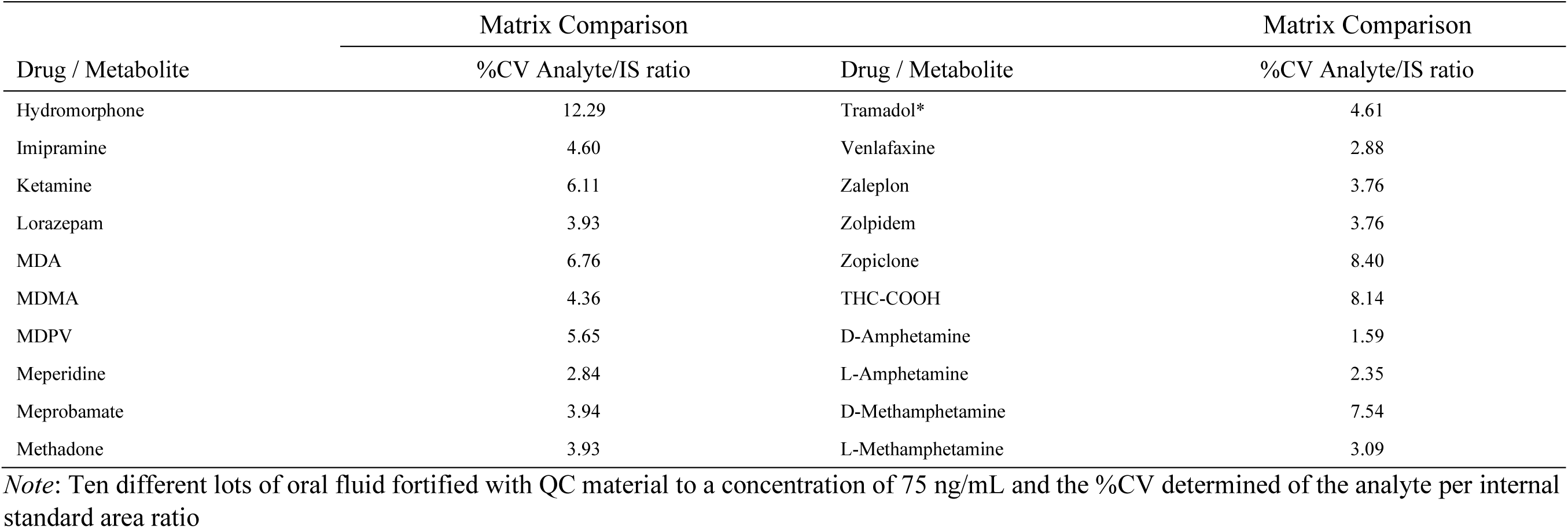
Matrix effects and recovery

### 3.5 Room temperature, refrigerator, and freezer stability

Samples with concentrations of 75 and 1800 ng/mL were prepared in triplicate. One set was kept at room temperature overnight (RT), a second set was kept in the refrigerator overnight (RF) and a third set was kept in the freezer overnight (FZ). These validation samples were then run and compared to a triplicate preparation of QC samples that had been analyzed as normal. All results show less than 20% deviation from expected (Table 9).

### 3.6 Freeze/thaw (FT) stability

Validation samples with concentrations of 75 and 1800 ng/mL were frozen at −20 °C and thawed in sequence with samples taken after each FT cycle for a maximum of three cycles. These validation samples were analyzed in triplicate and compared to a triplicate preparation of validation samples that had not been subjected to this freeze/thaw cycle. The experimental results showed all meeting acceptance criteria.

### 3.7 Extracted sample stability

A stability experiment was performed where samples were stored in the instrument (3 day) or refrigerator (7 day) and re-injected after 3 and 7 days. All samples were within 20% of the initial results except for dihydrocodeine which was back within 20% at day 7.

### 3.8 Stability in matrix

A series of triplicate samples were analyzed over 7 days for stability at room temperature, 4 °C and −20 °C. The results indicated that all analytes were stable for at least 7 days refrigerated and frozen. Most of the analytes were stable at room temperature except zopiclone (decreased) and ritalinic acid (increased) at day 7. A study of 30 days confirmed these characteristics for zopiclone and ritalinic acid. It also showed that chlordiazepoxide decreased after 14 days at room temperature as did methylphenidate. The decreased methylphenidate appeared to correspond with the increase in ritalinic acid concentration (metabolite of methylphenidate).

### 3.9 Matrix effects and recovery

Table 10 shows the effect of 10 different matrix lots tested by using a series of 75 ng/mL samples prepared in water, MPA and 10 different matrices. The results were acceptable with less than 20% CV across oral fluid, water and MPA meeting acceptance criteria. This is likely due to dilution in 1.5 mL Quantisal extraction buffer before extraction.

### 3.10 Selectivity

Multiple drugs that might have a potential for interfering with the assay analytes were run in the assay. Samples of 500 µL of 75 ng/mL QC were placed in a series of tubes to be run in triplicate. To the first set 50 µL of MeOH was added to act as the control. To the remaining tubes 50 µL of sample containing dextromethorphan, diphenhydramine, phenylephrine, salicylic acid, or combo (includes acetaminophen, caffeine, chlorpheniramine, ibuprofen, naproxen, and pseudoephedrine). These solutions were obtained from Cerilliant and were at a concentration of 1 mg/mL each except for the over the counter mix which was 100 µg/mL. Each solution was diluted to 20 µg/mL in methanol and this solution was used to spike samples as indicated above. Table 11 shows the results from this study. All samples met the acceptance criteria.

**TABLE 11.** Stability with concomitant medications

| Drug / Metabolite | % Diff from MEOH Spike |  |  |  |  |  |
| --- | --- | --- | --- | --- | --- | --- |
|  | Dextromethorphan | Phenylephrine | Diphenhydramine | Salicylic Acid | Phentermine | OTC Mix |
| 6-MAM | 5.39 | 2.41 | 3.68 | 1.56 | -8.41 | -7.78 |
| 7-Amino Clonazepam | 10.72 | 8.18 | 12.04 | 33.47 | -2.47 | -3.61 |
| Alprazolam | 9.27 | 2.99 | 7.65 | 14.10 | -1.38 | -2.03 |
| Amitriptyline | 3.11 | 2.96 | 8.50 | 7.57 | 0.10 | -1.86 |
| Amphetamine | -0.92 | -0.06 | -0.84 | 1.89 | -0.06 | 1.39 |
| a-OH Alprazolam | -0.83 | 2.54 | 0.06 | 4.41 | 4.71 | -11.52 |
| Benzoyllecgonine | -1.03 | 6.40 | 0.22 | 2.51 | 9.56 | 11.44 |
| Buprenorphine | 4.23 | 2.26 | -0.20 | -1.07 | -3.00 | -0.86 |
| Carisoprodol | -3.26 | -0.50 | 5.13 | 15.63 | 3.65 | -0.59 |
| Chlordiazepoxide | 0.96 | 0.34 | 2.35 | 2.07 | 0.34 | 0.33 |
| Clonazepam | 6.15 | 7.14 | 16.19 | 34.17 | 0.08 | 0.14 |
| Codeine | 13.56 | 14.51 | 6.73 | -15.03 | -0.35 | 6.84 |
| Cotinine | 1.22 | 3.48 | 3.92 | 2.92 | 3.17 | -2.21 |
| Cyclobenzaprine | 4.09 | -1.03 | 2.01 | 4.16 | -1.94 | -4.27 |
| Desalkylflurazepam | 8.09 | 7.14 | 10.06 | 38.05 | 5.65 | -3.29 |
| Desipramine | -1.45 | 2.43 | 0.43 | -3.89 | 0.31 | -3.13 |
| Diazepam | 1.13 | 3.37 | 2.48 | -5.65 | -1.78 | -2.52 |
| Dihydrocodeine | 0.62 | 0.79 | -7.21 | 3.27 | 3.78 | -1.00 |
| Doxepin | 0.23 | 4.39 | 2.31 | 0.83 | 1.58 | -0.64 |
| EDDP | 1.90 | 0.28 | 0.19 | -3.66 | 2.54 | -3.46 |
| Fentanyl | -1.40 | 1.54 | 3.41 | 0.29 | 7.39 | -2.49 |
| Fluoxetine | -1.69 | 4.26 | 2.23 | 2.50 | -0.35 | -1.47 |
| Gabapentin | -11.11 | -8.90 | -4.18 | -2.32 | 19.22 | -13.38 |
| Hydrocodone | 6.16 | -2.30 | 3.15 | -11.25 | 6.89 | 4.67 |

| % Diff from MEOH Spike |  |  |  |  |  |  |
| --- | --- | --- | --- | --- | --- | --- |
| Drug / Metabolite | Dextromethorphan | Phenylephrine | Diphenhydramine | Salicylic Acid | Phentermine | OTC Mix |
| Hydromorphone | 1.28 | 1.26 | 18.21 | -12.76 | -7.42 | -1.09 |
| Imipramine | -2.95 | -0.25 | 4.92 | 2.80 | 6.61 | 14.32 |
| Ketamine | -2.95 | -0.25 | 4.92 | 2.80 | 6.61 | -4.10 |
| Lorazepam | -0.88 | -3.20 | 7.82 | 16.67 | -1.02 | -3.45 |
| MDA | 4.59 | 5.12 | 7.04 | -0.89 | -7.19 | -6.86 |
| MDMA | 0.60 | 3.63 | -3.09 | -8.02 | -0.52 | 1.36 |
| MDPV | 3.95 | 3.01 | -0.89 | 1.69 | -1.13 | 0.14 |
| Meperidine | -1.77 | -0.99 | 1.10 | -0.57 | 4.74 | 1.55 |
| Meprobamate | 3.82 | 0.70 | 3.78 | 0.56 | 1.04 | 0.40 |
| Methadone | -2.63 | -0.39 | -2.11 | -6.01 | 0.73 | -1.11 |
| Methamphetamine | 6.15 | 3.95 | 6.10 | 2.67 | -2.29 | -1.34 |
| Methylphenidate | -1.50 | 0.49 | -0.28 | 0.67 | 3.21 | 1.19 |
| Morphine | 1.68 | 2.00 | 1.62 | -7.46 | 4.21 | 1.92 |
| Naloxone | -4.45 | -2.67 | -9.62 | -2.56 | -9.02 | 3.23 |
| Naltrexone | -0.33 | -0.66 | -1.65 | -2.72 | 5.19 | -6.07 |
| Nicotine | -0.98 | -0.95 | 6.01 | -3.25 | -1.76 | -3.54 |
| Norbuprenorphine | -0.49 | -4.07 | -0.15 | -0.22 | 0.10 | -1.60 |
| Nordiazepam | 6.90 | 6.95 | 15.20 | 34.26 | 3.76 | -2.96 |
| Norfentanyl | 7.69 | 15.16 | 7.72 | -11.99 | 14.82 | 1.37 |
| Norhydrocodone | 0.52 | 19.05 | 9.66 | -1.51 | 11.61 | -0.85 |
| Norketamine | 3.12 | -0.17 | 8.56 | -0.95 | -2.72 | -5.45 |
| Noroxycodone | -2.46 | -2.81 | 10.90 | 17.18 | -5.45 | -4.41 |
| Noroxymorphone | 4.29 | 0.50 | 3.46 | 15.05 | 1.84 | -6.40 |
| Nortriptyline | 2.86 | 1.00 | 3.95 | 1.59 | -1.52 | 4.69 |
| O-desmethyl Tramadol | -6.42 | 2.66 | -1.98 | 4.48 | 12.13 | -7.60 |
| Oxazepam | 0.63 | -0.74 | 1.86 | -2.35 | 1.21 | -1.92 |
| Oxycodone | 5.93 | 1.50 | -4.83 | -9.02 | -12.78 | 15.17 |
| Drug / Metabolite | Dextromethorphan | Phenylephrine | Diphenhydramine | Salicylic Acid | Phentermine | OTC Mix |
| Oxymorphone | 3.75 | -2.74 | -0.13 | -2.40 | 3.74 | -1.73 |
| Paroxetine | 1.54 | 3.91 | 6.26 | 0.71 | 2.68 | -5.17 |
| Phencyclidine | -4.40 | 2.95 | -5.07 | 0.72 | 8.75 | -6.18 |
| Pregabalin | 32.10 | 7.43 | -0.44 | 3.95 | -2.33 | 12.00 |
| Ritalinic Acid | 2.93 | 1.66 | 2.32 | 5.83 | 5.40 | 3.83 |
| Tapentadol | 1.61 | 4.28 | 5.77 | 0.29 | 8.08 | 0.35 |
| Temazepam | -0.73 | 1.88 | 2.39 | -3.55 | 0.39 | 1.35 |
| Tramadol | 4.16 | 2.43 | 3.01 | 2.80 | 2.19 | 1.42 |
| Venlafaxine | -2.84 | -1.08 | -3.37 | -2.58 | 4.23 | -3.12 |
| Zaleplon | 4.85 | 1.81 | 2.18 | 2.85 | -2.68 | -1.70 |
| Zolpidem | 1.03 | 0.03 | 0.77 | 1.05 | 1.59 | -2.96 |
| Zopiclone | -1.91 | -4.91 | -3.47 | -18.30 | -0.47 | -3.27 |
| THC-COOH | 0.66 | -5.77 | 1.05 | 7.18 | -3.65 | 10.25 |
| D-Amphetamine | -0.77 | -1.64 | -2.00 | 1.06 | -0.94 | -3.90 |
| L-Amphetamine | -1.97 | -0.41 | 2.07 | 1.30 | 3.21 | -0.54 |
| D-Methamphetamine | -6.26 | 5.53 | -6.10 | -3.81 | 7.45 | -5.96 |
| L-Methamphetamine | -2.10 | -0.24 | -0.08 | 0.71 | 7.73 | -0.12 |

*Note:* The indicated medications prepared in methanol were spiked into a QC 75 standard and measured. The data indicates percent difference from a QC standard spiked with blank methanol at the same volume as the drug standards.

## 4 DISCUSSION

The determination of prescription medications and illicit substances in oral fluids is one of the most non-invasive and easily observed sample collection methods. It provides a relatively simple and reliable means of sample collection coupled with a reduced chance of sample adulteration. Oral fluid also provides a viable alternative for measurement in patients that cannot provide an adequate urine sample volume such as catheterized patients. Analytes in oral fluid do not require a deconjugation step as in urine samples because the drugs are not glucuronidated for excretion in oral fluids and provides a lower cutoff than urine analysis. The only drawbacks of the oral fluid assay are that it has a shorter detection window and requires a more sensitive assay.

This paper presents a cost-effective means of analysis using older, less sensitive instruments (API SCIEX 4000) by using a liquid-liquid extraction method, concentration of the samples with a nitrogen dry-down, and a resuspension step. We validated the methods in accordance with FDA guidelines^24^ with an LOQ of 5 ng/mL for most of the analytes except for fentanyl and THC-COOH at 1 ng/mL and gabapentin and pregabalin at 25 ng/mL. The ULOQ was 2000 ng/mL with a few exceptions that required a lower ULOQ of 1000 ng/mL These were amitriptyline, amphetamine, chlordiazepoxide, ketamine, MDA, meperidine, methylphenidate, ritalinic acid, and tramadol.

## 5 CONCLUSION

The novelty of this study is a sensitive and low-cost method of ODT developed and validated for the determination of 64 drug analytes and the D- and L- isomers of amphetamine and methamphetamine from the same liquid-liquid extraction on an older model API 4000 mass spectrometer. Separation and detection are based on 3 LC-MS/MS injections. This method has an 8 minute run-time for most of the analytes, 1.5 minutes for the THC-COOH, and 10 minutes for the D- and L- isomeric separation for amphetamine and methamphetamine. The assay is quite sensitive for the majority of the analytes with a cutoff of 5 ng/mL except for fentanyl and THC-COOH at 1 ng/mL and gabapentin and pregabalin at 25 ng/mL, The assay also has good precision and accuracy and would add a valuable option to high throughput laboratories seeking a robust testing alternative to UDT methods and for medical providers seeking to achieve medication compliance^27^.

## Abbreviations

LLOQ: lower limit of quantitation
L: Levo
D: Dextro
6-MAM: 6-Monoacetylmorphine
THC: delta-9 tetrahydrocannabinol
THC-COOH: 11-Nor-9-carboxy-Δ9-tetrahydrocannabinol
EDDP: 2-Ethylidene- 1,5-dimethyl-3,3-diphenylpyrrolidine
ES: Electrospray
MDA: 3,4-Methylenedioxyamphetamine
MDMA: 3,4-Methylenedioxymethamphetamine
MDPV: Methylenedioxypyrovalerone
ODT: oral drug testing
mL: milliliter
µL: microliter
ng: nanogram
mm: millimeter
cm: centimeter
µm: micrometer
MPA: Mobile Phase A
MPB: Mobile Phase B
MPDL: D and L Mobile Phase
DCM: dichloromethane
QS: quantum satis
LC/MS: liquid chromatography mass spectrometry
SC: standard curve
QC: quality control
LC-MS/MS: triple quadrapole mass spectrometer
MeOH: methanol
MRM: multiple reaction monitoring
CAP: College of American Pathologists
SOP: standard operating procedure
IS: internal standard
LOD: limit of detection
ULOQ: upper limit of quantitation
RT: room temperature
RF: refrigerated
FZ: frozen
OTC: over the counter
CE: collision energy
GS1: ion source gas 1
GS2: ion source gas 2
DP: declustering potential
EP: entrance potential
CXP: collision cell exit potential
RT: retention time
CEM: channel electron multiplier
CAD: Collision Gas
ISV: Ion Spray Voltage
Q1: Quadrapole 1
Q3: Quadrapole 3
P63: 63 drug panel
SD: standard deviation,
%CV: percent coefficient of variation
%E: percent error
AMR: analytical measurement range
ExS: Extraction solution

## Appendix A

Transitions and retention times P63, THC, and D- and L- amphetamine and methamphetamine

Appendix A Transitions and retention times P63, THC, and D- and L- amphetamine and methamphetamine
| Drug / Metabolite | RT<br>(min) | Q1 | Q3 | DP | EP | CE | CXP |
| --- | --- | --- | --- | --- | --- | --- | --- |
| 6 MAM | 2.58 | 328.2 | 165.2 | 100 | 10 | 51 | 13 |
| 6 MAM 2 | 2.58 | 328.2 | 193.2 | 80 | 10 | 40 | 10 |
| Hydromorphone D-6 | 1.94 | 292.1 | 185.1 | 126 | 10 | 41 | 16 |
| 7-Aminoclonazepam D-4 | 3.17 | 290.1 | 121.1 | 50 | 10 | 46 | 4 |
| 7-Aminoclonazepam | 3.17 | 286.1 | 121.1 | 85 | 10 | 40 | 20 |
| 7-Aminoclonazepam 2 | 3.17 | 286.1 | 250.1 | 85 | 10 | 28.6 | 15 |
| α-OH-Alprazolam D-5 | 4.69 | 330.2 | 302.1 | 50 | 10 | 36 | 15 |
| α-OH-Alprazolam | 4.69 | 325.2 | 297.2 | 51 | 10 | 33 | 6 |
| Alprazolam D-5 | 4.85 | 314.2 | 286.1 | 50 | 10 | 46 | 4 |
| Alprazolam | 4.85 | 309.2 | 205.2 | 56 | 10 | 55 | 4 |
| Amitriptyline | 4.63 | 278.1 | 91 | 50 | 10 | 23 | 8 |
| Fentanyl D-5 | 4.25 | 342.2 | 105.1 | 46 | 10 | 53 | 4 |
| Amphetamine D-51 | 2.06 | 141.1 | 96.1 | 66 | 10 | 23 | 15.5 |
| Amphetamine | 2.09 | 136 | 91.1 | 41 | 10 | 21 | 6 |
| Amphetamine 2 | 2.09 | 136 | 119.1 | 41 | 10 | 12.5 | 21 |
| Benzoylecgonine D-8 | 3.12 | 298.3 | 171.1 | 41 | 10 | 25 | 4 |
| Benzoylecgonine | 3.12 | 290.2 | 105.1 | 41 | 10 | 39 | 4 |
| Benzoylecgonine 2 | 3.12 | 290.2 | 168.1 | 65 | 10 | 27 | 27 |
| Buprenorphine D-4 | 4.3 | 472.3 | 59.1 | 80 | 10 | 89 | 8 |
| Buprenorphine | 4.3 | 468.4 | 55.0 | 100 | 10 | 90 | 13 |
| Fentanyl D-5 | 4.25 | 342.2 | 105.1 | 46 | 10 | 53 | 4 |
| Carisoprodol 2 | 4.37 | 261.1 | 97.2 | 40 | 10 | 24 | 14 |
| Carisoprodol | 4.37 | 261.1 | 176.4 | 60 | 10 | 17 | 8 |
| Zolpidem D-6 | 3.92 | 314.3 | 235.3 | 100 | 10 | 45 | 13 |
| Chlordiazepoxide | 3.97 | 300.1 | 227.1 | 60 | 10 | 36 | 15 |
| Oxazepam D-5 | 4.64 | 292.3 | 246.2 | 70 | 10 | 32 | 6 |
| Clonazepam | 4.73 | 316.0 | 270.1 | 80 | 10 | 36 | 14 |
| Clonazepam 2 | 4.73 | 316.0 | 213.9 | 80 | 10 | 53 | 30 |
| Codeine D-6 | 2.41 | 306.2 | 165.2 | 100 | 10 | 36 | 10 |
| Codeine | 2.41 | 300.2 | 152.1 | 56 | 10 | 83 | 4 |
| Codeine Qual | 2.41 | 300.2 | 44.2 | 80 | 10 | 70 | 6 |
| Cotinine D-3 | 1.02 | 180.0 | 80.0 | 46 | 10 | 35 | 4 |
| Cotinine | 1.02 | 177.1 | 80.2 | 46 | 10 | 35 | 4 |
| Cyclobenzaprine D-3 | 4.57 | 279 | 215 | 17 | 10 | 52 | 12 |
| Cyclobenzaprine HI | 4.58 | 276.2 | 216.3 | 46 | 10 | 35 | 4 |
| Cyclobenzaprine | 4.58 | 276.2 | 216.5 | 46 | 10 | 35 | 4 |
| Desalkylflurazepam D-4 | 4.70 | 293.2 | 140 | 71 | 10 | 41 | 25 |
| Desalkylflurazepam | 4.71 | 289.1 | 140.1 | 71 | 10 | 41 | 25 |
| Imipramine D-3 | 4.57 | 284.1 | 89.1 | 110 | 10 | 32 | 10 |
| Desipramine | 4.54 | 267.1 | 72.0 | 46 | 10 | 41 | 4 |
| Diazepam D-5 | 5.00 | 290.1 | 198.1 | 80 | 10 | 40 | 15 |
| Diazepam | 5.01 | 285.2 | 193.2 | 80 | 10 | 40 | 15 |
| Diazepam 2 | 5.01 | 285.1 | 154.1 | 80 | 10 | 41 | 4 |
| dihydrocodeine D-6 | 2.29 | 308.2 | 202.1 | 46 | 10 | 41 | 4 |
| dihydrocodeine | 2.32 | 302.1 | 199.1 | 50 | 10 | 46 | 4 |
| Doxepin D-3 | 4.41 | 283.1 | 107 | 75 | 10 | 32 | 6 |
| Doxepin | 4.41 | 280.1 | 107 | 61 | 10 | 39 | 4 |
| EDDP D-3 | 4.53 | 281.1 | 234.1 | 61 | 10 | 39 | 4 |
| EDDP HI | 4.54 | 278.2 | 186.3 | 50 | 10 | 46 | 6 |
| EDDP | 4.54 | 278.2 | 186.5 | 50 | 10 | 46 | 6 |
| Fentanyl D-5 | 4.25 | 342.2 | 105.1 | 46 | 10 | 53 | 4 |
| Fentanyl | 4.25 | 337.2 | 188.6 | 51 | 10 | 29 | 4 |
| Fluoxetine D-6 | 4.47 | 316.1 | 154.2 | 45 | 10 | 12 | 15 |
| Fluoxetine HI | 4.47 | 310.1 | 44.3 | 50 | 10 | 40 | 10 |
| Fluoxetine | 4.47 | 310.1 | 148.2 | 50 | 10 | 11 | 10 |
| Gabapentin D-10 | 1.99 | 182.3 | 164.4 | 80 | 10 | 20 | 6 |
| Gabapentin | 2.03 | 172.1 | 154.1 | 22 | 10 | 22 | 13 |
| Oxymorphone D-3 | 1.76 | 305.2 | 230.2 | 50 | 10 | 46 | 4 |
| Hydrocodone | 2.77 | 300.2 | 199.1 | 150 | 10 | 41 | 7 |
| Hydromorphone D-6 | 1.94 | 292.1 | 185.1 | 126 | 10 | 41 | 16 |
| Hydromorphone | 1.95 | 286.1 | 185.1 | 56 | 10 | 39 | 4 |
| Hydromorphone Qual | 1.95 | 286.1 | 128.1 | 80 | 10 | 77 | 12 |
| Imipramine D-3 | 4.57 | 284.1 | 89.1 | 110 | 10 | 32 | 10 |
| Imipramine | 4.57 | 281.1 | 58 | 110 | 10 | 32 | 10 |
| Ketamine D-4 | 3.13 | 242.1 | 129.1 | 40 | 10 | 36 | 22 |
| Ketamine HI | 3.12 | 238.1 | 125 | 110 | 10 | 32 | 10 |
| Ketamine | 3.12 | 238.1 | 125.3 | 110 | 10 | 32 | 10 |
| Fentanyl D-5 | 4.25 | 342.2 | 105.1 | 46 | 10 | 53 | 4 |
| Lorazepam | 4.63 | 321.2 | 275.2 | 81 | 10 | 23 | 4 |
| MDA D-5 | 2.43 | 185.1 | 110.1 | 45 | 10 | 31 | 7 |
| MDA | 2.45 | 180.0 | 135.2 | 45 | 10 | 25 | 12 |
| MDA 1 | 2.45 | 180.0 | 163.4 | 45 | 10 | 12.5 | 11.5 |
| MDMA D-5 | 2.68 | 199.1 | 165.1 | 80 | 10 | 17 | 4 |
| MDMA | 2.69 | 194.1 | 163.5 | 75 | 10 | 17 | 10 |
| MDMA 2 | 2.69 | 194.1 | 133.1 | 75 | 10 | 28 | 23 |
| MDPV D-8 | 3.66 | 284.2 | 134.5 | 46 | 10 | 29 | 4 |
| MDPV HI | 3.66 | 276.2 | 126.1 | 40 | 10 | 37 | 20 |
| MDPV | 3.66 | 276.2 | 126.5 | 40 | 10 | 37 | 20 |
| Meperidine D-4 | 3.56 | 252.2 | 224.2 | 46 | 10 | 29 | 4 |
| Meperidine | 3.57 | 248.1 | 174 | 46 | 10 | 29 | 4 |
| Meprobamate D-7 | 3.48 | 226.1 | 165.1 | 41 | 10 | 13 | 4 |
| Meprobamate HI | 3.51 | 219.2 | 158.2 | 41 | 10 | 13 | 4 |
| Meprobamate | 3.51 | 219 | 97.08 | 41 | 10 | 13 | 4 |
| Methadone D-3 | 4.65 | 313.1 | 105.1 | 60 | 10 | 38 | 18 |
| Methadone HI | 4.66 | 310.2 | 105.2 | 60 | 10 | 25 | 6 |
| Methadone | 4.66 | 310.2 | 104.7 | 60 | 10 | 25 | 6 |
| Methamphetamine D-5 | 2.38 | 155.1 | 92.1 | 36 | 10 | 27 | 4 |
| Methamphetamine | 2.39 | 150.1 | 119.1 | 60 | 10 | 20 | 15 |
| Methylphenidate D-9 | 3.36 | 243.1 | 93.2 | 50 | 10 | 46 | 4 |
| Methylphenidate | 3.33 | 234.1 | 56.1 | 80 | 10 | 64 | 8 |
| Morphine D-6 | 1.62 | 292.1 | 152.3 | 80 | 10 | 80 | 13 |
| Morphine | 1.65 | 286.2 | 152.2 | 80 | 10 | 78 | 13 |
| Morphine 2 | 1.65 | 286.1 | 201.1 | 80 | 10 | 35 | 10 |
| Naloxone D-5 | 2.32 | 333.2 | 273.3 | 100 | 10 | 38 | 10 |
| Naloxone | 2.34 | 328.0 | 212.1 | 70 | 10 | 51 | 10 |
| Naltrexone D-3 | 2.63 | 345 | 58.2 | 105 | 10 | 58 | 10 |
| Naltrexone | 2.64 | 342.2 | 324.1 | 80 | 10 | 29 | 20 |
| Nicotine D-4 | 0.64 | 167.1 | 136.2 | 50 | 10 | 30 | 6 |
| Nicotine | 0.66 | 163.1 | 130.2 | 50 | 10 | 28 | 23 |
| Norbuprenorphine D-3 | 3.78 | 417.3 | 83.1 | 90 | 10 | 69 | 15 |
| Norbuprenorphine | 3.79 | 414.2 | 83.1 | 110 | 10 | 66 | 15 |
| Nordiazepam D-5 | 4.74 | 276.2 | 140.1 | 61 | 10 | 39 | 4 |
| Nordiazepam | 4.74 | 271.2 | 140.1 | 56 | 10 | 37 | 4 |
| Norfentanyl D-5 | 2.96 | 238.1 | 155.2 | 75 | 10 | 25.3 | 5.8 |
| Norfentanyl | 2.98 | 233.1 | 150.1 | 56 | 10 | 25 | 10 |
| Norfentanyl 2 | 2.98 | 233.1 | 84.1 | 56 | 10 | 26 | 15 |
| Norhydrocodone D-3 | 2.59 | 289.2 | 244.1 | 70 | 10 | 34 | 15 |
| Norhydrocodone | 2.60 | 286.2 | 199.1 | 85 | 10 | 38 | 10 |
| Norhydrocodone 2 | 2.60 | 286.2 | 128.1 | 85 | 10 | 72 | 23 |
| Norketamine D-4 | 2.88 | 228.1 | 129.1 | 48 | 10 | 31 | 6 |
| Norketamine HI | 2.90 | 224 | 125.2 | 46 | 10 | 33 | 9 |
| Norketamine | 2.90 | 224.0 | 125.5 | 46 | 10 | 33 | 9 |
| Fentanyl D-5 | 4.25 | 342.2 | 105.1 | 46 | 10 | 53 | 4 |
| Noroxycodone | 2.50 | 302.1 | 227.1 | 70 | 10 | 40 | 10 |
| Noroxycodone 2 | 2.50 | 302.1 | 198 | 70 | 10 | 60 | 31 |
| Oxymorphone D-3 | 1.76 | 305.2 | 230.2 | 50 | 10 | 46 | 4 |
| Noroxymorphone | 1.37 | 288.2 | 213.2 | 70 | 10 | 40 | 10 |
| Noroxymorphone 2 | 1.37 | 288.2 | 184.2 | 70 | 10 | 62 | 14 |
| Nortriptyline D-3 | 4.59 | 267.2 | 233.3 | 67 | 10 | 20 | 14 |
| Nortriptyline HI | 4.59 | 264.1 | 91 | 52 | 10 | 33 | 15 |
| Nortriptyline | 4.59 | 264.1 | 91.4 | 52 | 10 | 33 | 15 |
| Oxazepam D-5 | 4.64 | 292.3 | 246.2 | 70 | 10 | 32 | 6 |
| Oxazepam | 4.65 | 287.1 | 241.1 | 41 | 10 | 27 | 6 |
| Oxycodone D-6 | 2.63 | 322.1 | 247.1 | 50 | 10 | 46 | 4 |
| Oxycodone | 2.67 | 316.1 | 241.2 | 75 | 10 | 38 | 19 |
| Oxymorphone D-3 | 1.76 | 305.2 | 230.2 | 50 | 10 | 46 | 4 |
| Oxymorphone | 1.76 | 302.1 | 227.3 | 80 | 10 | 38 | 10 |
| Oxymorphone 2 | 1.76 | 302.1 | 198.1 | 80 | 10 | 58 | 10 |
| Fentanyl D-5 | 4.25 | 342.2 | 105.1 | 46 | 10 | 53 | 4 |
| Paroxetine HI | 4.54 | 330.2 | 70.1 | 75 | 10 | 48 | 3.5 |
| Paroxetine | 4.54 | 330.2 | 70.5 | 75 | 10 | 48 | 3.5 |
| Phencyclidine D-5 | 4.25 | 249.1 | 86.1 | 15 | 10 | 17 | 10 |
| Phencyclidine HI | 4.26 | 244.1 | 91.2 | 45 | 10 | 40 | 15 |
| Phencyclidine | 4.26 | 244.1 | 91.6 | 45 | 10 | 40 | 15 |
| Pregabalin D-6 | 1.73 | 166.2 | 130.2 | 30 | 10 | 21 | 21 |
| Pregabalin | 1.75 | 160.0 | 55. | 30 | 10 | 34 | 10 |
| Pregabalin 2 | 1.75 | 160.0 | 83.1 | 30 | 10 | 35 | 15 |
| Phencyclidine D-5 | 4.25 | 249.1 | 86.1 | 15 | 10 | 17 | 10 |
| Ritalinic Acid | 2.88 | 220.1 | 84.1 | 95 | 10 | 22 | 10 |
| Tapentadol D-3 | 3.29 | 225.1 | 107 | 95 | 10 | 31 | 10 |
| Tapentadol HI | 3.28 | 222.1 | 121 | 95 | 10 | 31 | 10 |
| Tapentadol | 3.28 | 222.1 | 121.5 | 95 | 10 | 31 | 10 |
| Temezepam D-5 | 4.86 | 306.1 | 288 | 55 | 10 | 20 | 16 |
| Temezepam | 4.87 | 301.2 | 255.3 | 31 | 10 | 27 | 4 |
| Tramadol D-3 C-13 | 3.20 | 268.2 | 58.1 | 20 | 10 | 30 | 15 |
| Tramadol HI | 3.21 | 264.1 | 58.2 | 20 | 10 | 30 | 15 |
| Tramadol | 3.21 | 264.1 | 58.5 | 20 | 10 | 30 | 15 |
| O-desmethyl Tramadol HI | 2.44 | 250.1 | 58.3 | 20 | 10 | 30 | 15 |
| O-desmethyl Tramadol | 2.44 | 250.1 | 57.6 | 20 | 10 | 30 | 15 |
| O-desmethyl Tramadol D-6 | 2.44 | 256.2 | 64.2 | 20 | 10 | 30 | 15 |
| Venlafaxine D-6 | 3.75 | 284.2 | 64.2 | 50 | 10 | 43 | 10 |
| Venlafaxine HI | 3.75 | 278.2 | 57.7 | 50 | 10 | 41 | 11 |
| Venlafaxine | 3.75 | 278.2 | 57.6 | 90 | 10 | 23 | 11 |
| Zaleplon D-4 | 4.69 | 310.2 | 240.2 | 90 | 10 | 38 | 17 |
| Zaleplon | 4.69 | 306.2 | 236.3 | 85 | 10 | 37 | 7 |
| Zolpidem D-6 | 3.92 | 314.3 | 235.3 | 100 | 10 | 45 | 13 |
| Zolpidem | 3.94 | 308.1 | 235.2 | 20 | 10 | 30 | 15 |
| Oxymorphone D-3 | 1.76 | 305.2 | 230.2 | 50 | 10 | 46 | 4 |
| Zopiclone | 3.56 | 389.1 | 244.8 | 20 | 10 | 30 | 15 |

Transitions and retention times for THC

| Drug Name | Retention<br>time | Parent >daughter(m/z) | DP | EP | CE | CXP |
| --- | --- | --- | --- | --- | --- | --- |
| THC-COOH | 0.89 | 343.2>245.1 | -85 | -10 | -38 | -7 |
| THC-COOH-D9 | 0.87 | 352.2>308.2 | -85 | -10 | -38 | -7 |

Transitions and retention times for D- and L- amphetamine and methamphetamine

| Drug | RT<br>(min) | Q1 | Q3 | DP | EP | CE | CXP |
| --- | --- | --- | --- | --- | --- | --- | --- |
| D-Amphetamine (S) | 6.50 | 136 | 91.000 | 35 | 10 | 25 | 6 |
| L-Amphetamine (R) | 7.21 | 136 | 91.101 | 35 | 10 | 25 | 6 |
| D Amphetamine D-11 | 6.51 | 147.2 | 98.100 | 35 | 10 | 26 | 7 |
| L Amphetamine D-11 | 7.22 | 147.2 | 98.101 | 35 | 10 | 26 | 7 |
| D-Methamphetamine (S) | 7.91 | 150 | 119.100 | 10 | 10 | 25 | 9 |
| L-Methamphetamine R | 8.45 | 150 | 119.101 | 10 | 10 | 25 | 9 |
| D Methamphetamine D-11 | 7.91 | 161 | 97.100 | 85 | 10 | 26 | 7 |
| L Methamphetamine D-11 | 8.46 | 161 | 97.101 | 85 | 10 | 26 | 7 |

*Note:* Retention times are updated using the internal standard each time that the data are analyzed.

